# Phylogenetic-based methods for fine-scale classification of PRRSV-2 ORF5 sequences: a comparison of their robustness and reproducibility

**DOI:** 10.1101/2024.05.13.593920

**Authors:** Kimberly VanderWaal, Nakarin Pamornchainvakul, Mariana Kikuti, Daniel Linhares, Giovani Trevisan, Jianqiang Zhang, Tavis K. Anderson, Michael Zeller, Stephanie Rossow, Derald J. Holtkamp, Dennis N. Makau, Cesar A. Corzo, Igor A.D. Paploski

## Abstract

Disease management and epidemiological investigations of porcine reproductive and respiratory syndrome virus-type 2 (PRRSV-2) often rely on grouping together highly related sequences. In the USA, the last five years have seen a major paradigm shift within the swine industry when classifying PRRSV-2, beginning to move away from RFLP (restriction fragment length polymorphisms)-typing and adopting the use of phylogenetic lineage-based classification. However, lineages and sub-lineages are large and genetically diverse, and the rapid mutation rate of PRRSV coupled with the global prevalence of the disease has made it challenging to identify new and emerging variants. Thus, within the lineage system, a dynamic fine-scale classification scheme is needed to provide better resolution on the relatedness of PRRSV-2 viruses to inform disease management and monitoring efforts and facilitate research and communication surrounding circulating PRRSV viruses. Here, we compare potential fine-scale systems for classifying PRRSV-2 variants (i.e., genetic clusters of closely related ORF5 sequences at finer scales than sub-lineage) using a database of 28,730 sequences from 2010 to 2021, representing >55% of the U.S. pig population. In total, we compared 140 approaches that differed in their tree-building method, criteria, and thresholds for defining variants within phylogenetic trees using *TreeCluster*.

Three approaches produced epidemiologically meaningful variants (i.e., ≥5 sequences per cluster), and resulted in reproducible and robust outputs even when the input data or input phylogenies were changed. In the three best performing approaches, the average genetic distance amongst sequences belonging to the same variant was 2.1 – 2.5%, and the genetic divergence between variants was 2.5-2.7%. Machine learning classification algorithms were also trained to assign new sequences to an existing variant with >95% accuracy, which shows that newly generated sequences could be assigned without repeating the phylogenetic and clustering analyses. Finally, we identified 73 sequence-clusters (dated <1 year apart with close phylogenetic relatedness) associated with circulation events on single farms. The percent of farm sequence-clusters with an ID change was 6.5-8.7% for our best approaches. In contrast, ∼43% of farm sequence-clusters had variation in their RFLP-type, further demonstrating how our proposed fine-scale classification system addresses shortcomings of RFLP-typing. Through identifying robust and reproducible classification approaches for PRRSV-2, this work lays the foundation for a fine-scale system that would more reliably group related field viruses and provide better improved clarity for decision-making surrounding disease management.

## 1. Introduction

In infectious disease epidemiology, the ability to identify closely related genetic sequences is important for epidemiological investigations and tracking pathogen spread. There is no standard or universal approach for classifying and naming viral genetic diversity below the species level. However, labeling genetic clusters facilitates monitoring, research, and communication regarding viral genetic diversity present in host populations (1). In addition, the ability to cluster sequences according to genetic relatedness lays the foundation for research evaluating phenotypic variation amongst different named clades of the virus.

These applications of sequencing are particularly relevant for porcine reproductive and respiratory syndrome virus (PRRSV), where sequencing is not only used for research purposes, but also for day-to-day disease management implemented by animal health professionals in the field (2, 3). In the U.S., PRRSV is primarily caused by PRRS virus-type 2 (PRRSV-2), a positive-sense single-stranded RNA virus (the species *Betaarterivirus suid 2* in the genus *Betaarterivirus*, family *Arteriviridae*, order *Nidovirales*) (3, 4). With an economic burden of >$600 mill/year in the U.S. (5), PRRSV-2 is the most important endemic disease in the U.S. swine industry (6, 7), impacting farrowing rates, number of weaned pigs, poor growth, and mortality (5, 8). Approximately 30-50% of U.S. breeding farms have active PRRSV circulation within their herds (9, 10). The rapid evolution and genetic and antigenic diversity of PRRSV-2 are key complicating factors to the control of this disease (3, 11–13). In addition to the co-circulation of numerous lineages and sub-lineages of the virus, the routine emergence of novel sub-lineages creates recurring epidemic waves that spread rapidly and widely through the industry (14–16).

Sequencing is used as part of routine disease monitoring on farms, primarily to discriminate newly introduced viruses from previous/resident strains present on a farm, determine the possible origin of between-farm spread, and inform choice of immunization strategy (17). Within the ∼15 kb PRRSV-2 genome, open reading frame 5 (ORF5) encodes for a major envelope protein (glycoprotein 5 - GP5), which is involved in inducing virus neutralizing antibodies and cross-protection among PRRSV variants (18–20). Even though ORF5 accounts for only 4% of the genome, its genetic variability and apparent immunologic importance (3, 20–22) has made this gene the target of nearly all genetic sequencing conducted by the swine industry, with thousands of sequences generated per year in the U.S. alone (4). Stakeholder preference for ORF5 rather than whole genome sequencing also relates to lower cost, rapid turnaround time, and the higher probability of successfully obtaining a sequence from samples of various types and quality.

While phylogenetic analysis is the gold standard for interpretation of sequence data, animal health professionals in the field often find it timelier and more convenient to have a name that they can use to refer to a given cluster of genetic sequences as part of everyday communication and outbreak investigations. Additionally, most sequences are not publicly available, so phylogenetic analysis can potentially lead to different interpretations if the available data for practitioners are not representative. Thus, relying on universal classifications assures standardization between different users. Currently, the naming method used by the industry to discriminate between sequences is RFLP-typing (Restriction Fragment Length Polymorphisms), sometimes in combination with an additional label corresponding to phylogenetic lineage (3, 14, 15, 23, 24). However, using RFLP-types to refer to PRRSV-2 viruses often leads to misleading or even erroneous conclusions (e.g., viruses assigned to the same RFLP-type often are not closely related, and vice versa)(15, 25). Moreover, only 6 sub-lineages are currently prevalent in the U.S. (at least 5% of detected sequences from 2019-23, (26)), each typically having a mean genetic distance of <8.5% (but occasionally higher than 10%) for sequences belonging to the sub-lineage (23), making these classifications too coarse for on-farm disease monitoring and decision-making.

Thus, better methods are needed to further sub-divide the wide genetic diversity present within lineages and sub-lineages into smaller groups of closely related sequences (termed “variants”) that facilitate monitoring, research, and communication surrounding genetic diversity. Fine-scale phylogenetic classification of PRRSV-2 is hampered by several challenges common to most RNA viruses. Building phylogenetic trees with thousands of sequences is increasingly common in studies of viral evolution and molecular epidemiology, and instability in topology is often an issue when building such large phylogenies based on sequences that are closely related and relatively short in length (i.e., using marker genes such as ORF5 for PRRSV-2)(27). Thus, while partitioning trees into clusters that meet user-set criteria (such as the maximum genetic distance allowable within a cluster) is relatively straightforward using existing methods (28, 29), it is less clear which methods and their associated user-set criteria produce stable clusters that are robust and reproducible when analyses are repeated with different sets of data. However, such reproducibility is essential for any classification system.

The purpose of this paper is to evaluate different phylogenetic clustering approaches that could be used as the basis of a fine-scale classification of PRRSV-2 in the U.S. Particularly, we aim to identify methodology that would overcome the shortcomings of the current PRRSV-2 nomenclature systems, and that may be reliable for the analysis of large phylogenies of RNA viruses more generally. Due to the paucity of whole genome sequence data and severe limitations in our understanding of genotype-phenotype interactions, we note that existing PRRSV-2 classification systems and the additional refinements explored here are not meant to be based on phenotypic variation of the virus, but rather have application for epidemiological monitoring.

## 2. Methods

### 2.1 Data Source and sequence alignment

Sequence data were obtained from the Morrison Swine Health Monitoring Project (MSHMP), which is a voluntary initiative operated by University of Minnesota that monitors PRRS occurrence in the U.S. MSHMP was initiated in 2011, and currently collects weekly infection status data for breeding farms belonging to 37 production systems, accounting for >55% of the U.S. sow population (10). Participating production systems also share PRRSV ORF5 sequences that are generated as part of routine monitoring and outbreak investigations in breeding, gilt developing units, growing and finishing herds (4). Sequences are generally obtained either directly from each MSHMP participant or from the main veterinary diagnostic laboratory where participants submit their diagnostic samples. Meta-data for each sequence include farm ID (anonymized), sample collection date, and farm type of origin (e.g., breeding or growing herd). Sequences without a farm ID or location information were excluded.

Sequences were divided into short-and long-term datasets. The short-term dataset, which included three years of sequence data (6,749 sequences from July 1, 2018 – June 30, 2021), was utilized for developing and comparing different classification methods in classifying PRRSV-2 genetic variants that co-circulate within U.S. swine populations. The long-term dataset, which included ∼11 years of sequence data (28,965 sequences from January 1, 2010 – September 30, 2021) was used to evaluate the farm-level occurrence of PRRSV variants.

### 2.2 Tree building

Sequence datasets were cleaned to exclude ORF5 sequences with fewer than 603 bases or with more than 4 (0.5%) ambiguous bases. This resulted in 6646 sequences for the short-term dataset, and 28,730 sequences for the long-term dataset. Sequences were then aligned using the MAFFT’s local pairwise alignment algorithm (30, 31). Following this, de-duplicated datasets (n=4502 and n=13,721 for the short- and long-term datasets, respectively) were generated by eliminating sequences with 100% nucleotide identity.

All tree-building utilized in this analysis was performed using IQ-TREE with 1,000 ultrafast bootstraps (32, 33). Substitution model selection was performed for the short-term dataset using IQ-TREE, and we selected the model with the lowest BIC that was also widely available in other phylogenetic software platforms (to facilitate reproducibility of genetic variants based on phylogenetic clustering). Thus, the general time reversible substitution model with empirical base frequencies and gamma plus invariant site heterogeneity (GTR+F+I+G4) was selected and used for all subsequent tree-building described herein. Three tree types were generated for each dataset: a) a maximum-likelihood tree, b) a strict majority-rule consensus tree from the bootstrap trees (-minsup = 0.5, clades are collapsed if bootstrap support is < 0.5), and c) an extended majority-rule consensus tree (-minsup = 0). The *ggtree* package in R was used for all tree visualizations, with trees re-rooted on Lineage 5, which contains the PRRSV-2 prototype virus (11, 34).

### 2.3 Variant classification

A tree-based clustering approach was applied to the phylogenies using the *TreeCluster* package available in Python (28); clusters of genetically related sequences identified in the trees were referred to as “variants.” Briefly, we used six different methods available within this package to identify clusters of sequences within a tree: *Average clade (AC):* the average pairwise patristic distance between sequences within a variant is no more than *x*; the cluster must also form a monophyletic clade (i.e., include all descendent sequences from the clusters common ancestor). *Median clade (Med):* the median pairwise patristic distance between sequences within a variant is no more than *x*; the cluster must also form a monophyletic clade. *Length clade (LenC):* a variant does not contain any branches that are greater than length *x*; the cluster must also form a monophyletic clade. *Length (Len)*: same as LenC, but the variant need not form a monophyletic clade. *Single linkage (SL):* the distance between any two sequences in the variant cannot exceed *x*; the variant need not form a monophyletic clade. The SL method is analogous to the distance-based snowball method used previously to identify PRRSV-2 sequences involved in particular outbreaks (15). For all methods, thresholds (*x*) values of 2, 3, 4, and 5% were used. Early exploration suggested that the average clade method at the upper threshold of 5% produced clusters that were visually well aligned to phylogenetic structure in the tree; thus, higher thresholds of 6, 7, and 8% were also considered for the average clade method to assess if higher thresholds produced improved results. *TreeCluster’*s “support” argument specifies that the branches connecting every pair of sequences within a cluster must exceed a user-specified bootstrap support value; this was set to 0, given that sequences within a cluster are highly related and thus topological uncertainties internally within the cluster result in low bootstrap values. This is a particular issue when dealing with large sequence datasets for RNA viruses (27), and setting a higher support value results in clades becoming overly granular. For our purposes, it was more important for the ancestral node of the clade to have high bootstrap support, thus supporting the existence of the clade overall regardless of its exact internal topology. While *TreeCluster* does not evaluate support at the ancestral node, we summarized this value as part of our analysis below. For comparison, we also grouped sequences according to their RFLP-type as well as the combination of (sub-)lineage+RFLP (Lineage classification was used for lineages 2 – 9, and Sub-lineage was used to further stratify Lineage 1, which accounted for >70% of the sequences). In total, 140 approaches were compared: 23 *TreeCluster* methods applied to each of three tree types (maximum-likelihood, strict consensus, and extended consensus) built on two datasets (full and de-duplicated), plus RFLP and Lineage+RFLP. For the de-duplicated analysis, clustering was determined on the de-duplicated trees so that identical sequences did not pull down mean and median patristic distances within the *TreeCluster* analysis. The duplicate sequences were assigned to the same ID after running *TreeCluster*, and initial genetic characterization included these duplicate sequences.

### 2.4 Initial genetic characterization

For the short-term dataset, initial characterization of variants produced by each approach included a) the number of variants identified, b) the number of “common” variants (n >50 sequences belonging to the variant), c) the number of “singleton” variants, d) median sequences per variant and interquartile range, e) percent of sequences belonging to common variants, f) percent of sequences belonging to rare variants (n <10 sequences), g) median bootstrap value of the ancestral node, and h) mean genetic distance (raw p-distance) within a variant. Taking the within-variant means, we also summarized the i) the 95^th^ percentile of means across variants. Finally, we calculated the j) minimum genetic distance from each variant to the most closely related variant.

For subsequent analysis, we included only approaches that produced variants with a median of >5 sequences per variant and no less than 15% of sequences belonging to rare variants. Only 31 approaches met the criteria, which were subsequently compared to RFLP and Lineage+RFLP.

### 2.5 Reproducibility of classification amongst sets of data

Given that it is important for any proposed classification scheme to produce consistent results when applied to PRRSV-2 trees based on different data, we performed several analyses to determine the extent to which the variant classification produced above could be replicated when the data was re-analyzed. Trees utilized to assess the reproducibility of classification schemes included: a) a tree based on a duplicate IQ-TREE run utilizing the same data as above; b) a longer-term tree focused on one sub-lineage; c) a variant associated with a regional PRRSV-2 outbreak defined *a priori* from a previous analysis; d) subsets of data; and e) time-scaled Bayesian trees, which are often considered the gold-standard phylogenetic reconstruction.

a) *Duplicate tree*: For this reproducibility analysis, a duplicate phylogenetic tree was generated by running the same dataset in IQ-TREE with the same settings and different random seeds, as different runs of IQ-TREE can produce different trees due to the underlying stochastic algorithm used to find the tree with maximum likelihood (35). Variant classification was performed as described above, and the resulting classification (scheme B) was compared to the original classification (scheme A). The concordance between the classifications produced for each tree was quantified through the Jaccard index. Essentially, the Jaccard index quantifies how often pairs of sequences are assigned to the same variant ID across both schemes. It is calculated as the number pairs that are assigned to the same variant in scheme A and B (A=B) divided by the number of pairs that are assigned to the same variant in scheme A but not B and vice versa. The Jaccard index ranges between 0 and 1, with 1 indicating perfect concordance between scheme A and B.

b) *Sub-lineage tree with 15 years of data:* For this reproducibility analysis, 7,067 sequences for sub-lineage 1C from a time period of November 27, 2007 – November 21, 2022 were used, which included 797 of the same sequences present in our short-term dataset (all L1C sequences in the short-term dataset were also in this L1C dataset). This data set allowed us to ascertain the extent to which variant classifications produced for these two datasets differed when considering longer timeframes and more sequences from a single sub-lineage (i.e., the genetic diversity within this sub-lineage was more densely sampled). The Jaccard index was used to quantify the concordance between the classifications produced from each tree. Because not all sequences in one dataset appeared in the other, the Jaccard index was only calculated from sequence pairs that were present in both sets.

c) A priori *defined outbreak variant:* Since 2020, the swine industry in the Midwestern U.S. has witnessed large-scale spread of a novel PRRSV-2 variant, denoted as either L1C.5 or alternatively L1C-1-4-4 variant based on its sub-lineage and RFLP pattern (15, 23). A previous study used the inclusion criteria of >98% nucleotide identity to any of other sequences to define this clade. While this clade has a distinct recombination profile in genomic regions outside of the ORF5 gene, viruses belonging to this variant have >98% nucleotide identity at the whole genome level and form a distinct monophyletic clade within phylogenetic trees based on whole genomes or the ORF5 gene (36). This clade does not receive a distinguishing label in lineage classification, RFLP-typing, or the combination of both. To determine whether our classification methods capture this clade, we calculated the Jaccard index between our variant classification schemes and the L1C-1-4-4 clade as *a priori* defined by Kikuti et al (15).

d) *Trees based on 10 different subsets of the dataset:* To ascertain whether the classifications defined on the full short-term tree are robust across different subsets of data, we created 10 maximum-likelihood trees based on a distinct 10% of the short-term dataset– partitioning of sequences to sub-sets was random. Ideally, sequences that were classified together on the full tree should remain clustered on the subset trees. To assess this, we calculated two measures: clade purity and nearest neighbor matching. Clade purity was calculated for each variant *j* in a sub-tree by first identifying the clade containing those sequences by finding the most recent common ancestor (MRCAj) of all sequences belonging to that variant. We then identified all sequences descending from that ancestral node. Ideally, the descendent clade should purely contain sequences belonging to that variant; if sequences belonging to other variants were present within the descendent clade, this would indicate instability in the variant classification when constructing trees from smaller sets of data. We quantified the extent to which sequences belonging to other variants were present in variant *j*’s clade by calculating clade purity (proportion of sequences descending from the MRCA that belong to variant *j,* with 1 indicating perfect purity).

Clade purity was highly sensitive to single outlier sequences; if a single sequence is placed far away from the rest of the variant, then this results in a deep node being identified as the common ancestor, which means that a very large number of non-variant sequences are included in the clade, resulting in low purity metrics that are driven by a single outlier. To overcome the disproportionate effect of outlier sequences, we also performed nearest neighbor matching. Here, we identified the nearest neighbor for every sequence, and then tabulated whether the nearest neighbor belonged to the same variant or a different variant. We then calculated the proportion of sequences whose nearest neighbors were a member of the same variant. For both metrics, the median and interquartile of each metric across all sub-trees was reported. Only variants that had >1 sequence present in the sub-tree were considered.

e) *Trees constructed with BEAST:* Given computational constraints, time-scaled trees were constructed separately for each sub-lineage using BEAST v.1.10.4 (75 sequences for L1B, 568 for L1C, 795 sequences for L1H, and 129 for L1E sequences (23)). The temporal signal and approximate time to the most recent common ancestor (tMRCA) of each sub-lineage was first estimated from maximum-likelihood trees using TempEst (37). The models were run with the GTR, gamma distributed nucleotide substitution model, a lognormal uncorrelated relaxed molecular clock, and the Bayesian Skygrid population coalescence model (37) with 50 parameters and the time at last transition point set to the estimated tMRCA from TempEst (or 10 years if the estimated tMRCA was <10 years). Duplicate MCMC chains of 100 million steps each were run, sampling every 10,000^th^ step. For L1A (the largest sub-lineage with 1675 sequences), sequences were down-sampled to 777 sequences using an in-house script that maintains genetic and temporal diversity (i.e., the most closely related clades - i.e., clades where all branches have a length of ≤1 base - are first down-sampled to include only a single sequence per calendar quarter, followed iteratively by down-sampling clades with branch lengths less than 2 and then 3 bases, and so forth until no more than 800 sequences remained). In addition, only 25 Skygrid parameters were used for the L1A analysis to reduce computational burden and a chain length of 300 million was required for convergence. The first 10% of MCMC chains were discarded as burn-in. The duplicate runs were inspected for convergence, and then combined using LogCombiner. Maximum clade credibility trees (MCC) were built using TreeAnnotator v.1.10.4 and visualized with *ggtree*.

### 2.6 Ease of classification

For prospective application of any classification system, it is desirable to be able to classify new sequences to their respective variants without performing computationally heavy analysis. While acknowledging that any algorithm to classify sequences would need to be routinely updated as the virus evolves, we at the very least wanted to ensure that there were viable and accurate algorithms that could discriminate the fine-scale genetic variants defined in this paper. Therefore, we trained a random forest machine learning algorithm to assign new sequences to the appropriate variant grouping. This is the method used for variant assignment for SARS-CoV-2 (38). The model was trained using the first 90% of the short-term dataset (i.e., the ‘training’ dataset), holding out 10% of the most temporally recent data for model validation. Using the training dataset, the random forest was fitted using the *caret* package in R using ten-fold cross-validation and auto-tuning of the *mtry* hyper-parameter (39).

Model performance on the training set was assessed using ten-fold cross-validation (i.e., performance evaluated on 10% of observations that were left out of 10 iterative random forest runs). Two testing datasets were used: A) *internal test data:* the remaining (and most recent) 10% of the short-term dataset to simulate assignment of new sequences generated prospectively, and B) *external test data:* 4661 sequences from the University of Minnesota Veterinary Diagnostic Laboratory from the same three year period (only sequences that were <100% nucleotide identity to training sequences were used for model testing – 55.6% of sequences had 100% identity with the training dataset since these two datasets overlap). We report the overall accuracy (percent of sequences correctly classified by the algorithm) for the training dataset and for testing set A. We also calculated the mean groupwise precision, recall, and accuracy (i.e., percent of sequences correctly classified per variant was first calculated, and then a mean of these groupwise accuracies was reported). The true variant IDs were not known for testing set B given that these sequences were not part of the original variant classification analysis. Therefore, to assess the accuracy of assignments, we constructed a phylogenetic tree for test set B, and assigned the sequences that had 100% nucleotide identity with training sequences to their corresponding variant. We then performed nearest neighbor matching, as described above, and reported the proportion of testing sequences (with predicted variant IDs) whose nearest training sequence (with known variant IDs) in the tree belonged to the same variant. Only variants with more than one representative on the tree were considered in the calculation.

### 2.7 Farm-level occurrence of variants

We used the MSHMP database to tabulate the number of farms (based on their unique premises ID), number of production systems, and number of U.S. states (median and interquartile range) in which each variant was detected. Summaries were generated excluding rare variants (<10 sequences).

We also analyzed data for any farm in which ≥4 sequences in a single year were available to assess the stability of variant classification during micro-evolution that may occur during the course of an on-farm outbreak. 73 farms met this criterion, from which 587 sequences were available between January 12, 2010 and September 7, 2021 (4 – 43 sequences per farm, with some farms meeting these criteria in multiple years). Sequences were assigned to variant ID based on the classification methods applied to the long-term dataset. Here, we focused on the average clade (ac).06, ac.07, ac.08 methods from the strict consensus tree constructed with de-duplicated data, as this approach yielded the best results in the above analyses. A time-scaled tree was built for the 587 farm sequences using the same settings in the above BEAST analysis. For each classification method, we measured the maximum genetic distance and maximum divergence time (branch lengths represent time in a time-scaled phylogenetic tree) across every pair of sequences belonging to the same variant on the same farm. For each pair of variants that occurred on the same farm, we also measured the maximum genetic distance and maximum divergence time between sequences belonging to those variants. This enabled us to flag situations where sequences that were assigned to two variants actually formed a tight “sequence-cluster” for that farm in the phylogenetic tree (i.e., short genetic distances and divergence time between the two variants on the farm, thus likely representing the circulation of a single variant on the farm, Figure 1). Merging thresholds were established based on maximum genetic distance and divergence time to identify sequences that would be more accurately represented by a single variant ID rather than two IDs (i.e., they cluster together in the tree with short divergence times, hence they likely represent a farm sequence-cluster associated with a single circulation event).

The percent of farm sequence-clusters with ID changes was calculated as the number of variant-pairs that met the merging threshold (i.e., a farm sequence-cluster that has two associated IDs, see Figure 1) divided by the total number of farm sequence-clusters (i.e., farm sequence-clusters represented by a single ID detected). Of note, merging thresholds were applied only to quantify how often ID-changes occurred, but were not applied to the overall classification outlined in this paper.

**Figure 1.**
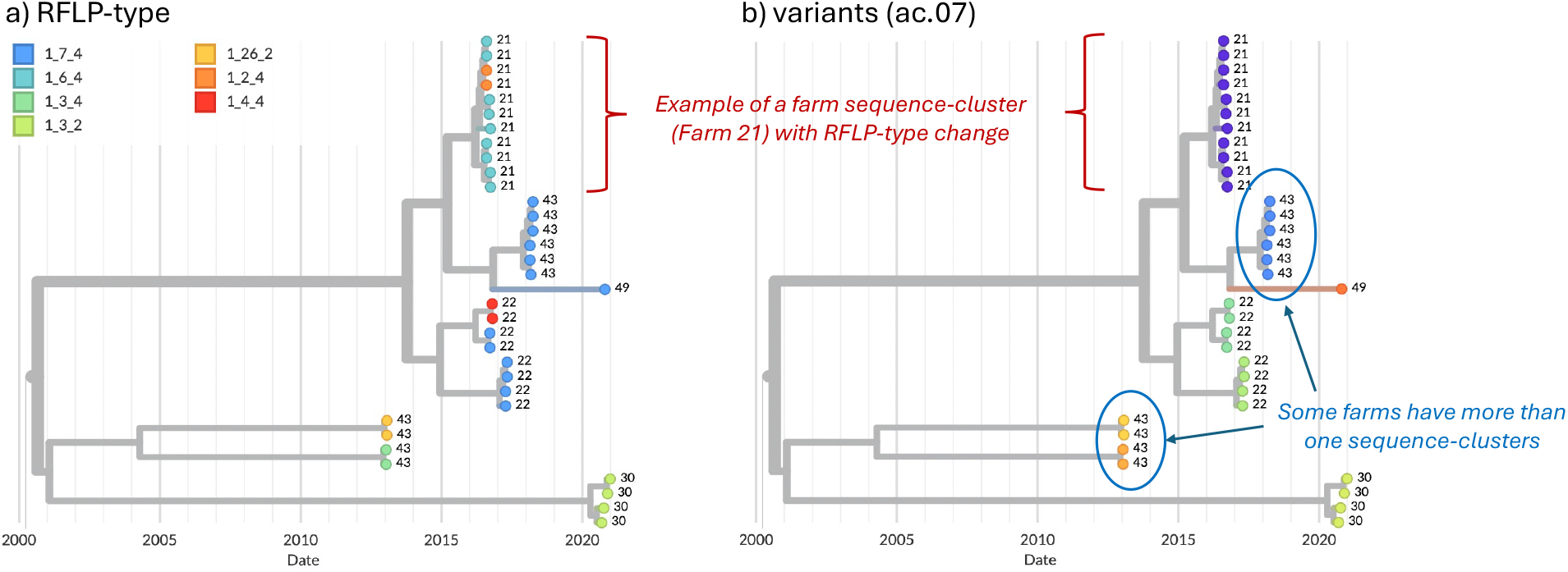
Example of farm sequence-clusters from five farms, visualized in Nextstrain (40). Farm ID is shown as the tip-labels. a) Tip colors represent RFLP-type. b) Tip colors represent the variant ID (ac.07 method).

## 3. RESULTS

### 3.1 Initial characterization of variants

Of the 140 classification approaches initially considered, 31 approaches met the initial criteria of having a median >5 sequences and no more than 15% of sequences belonging to “rare” variants (i.e., fewer than 10 sequences/variant). Only approaches that used a patristic distance threshold of ≥4% met this criterion, and 30 out of 31 approaches utilized the Average Clade method (denoted as ac.04, ac.05, etc., with the latter digits representing the patristic distance cutoff). These 31 candidates were compared to classifications based on RFLP-typing and Lineage+RFLPs, for a total of 33 approaches. Summary metrics for the average clade (ac) method applied to the strict consensus, deduplicated (con50.dedup) trees are shown in Table 1. Summary metrics for all approaches are shown in Supplementary Table S1.

As outlined in detail below, the overall best approaches (highlighted in green in Figure 3) were selected from amongst these 31 candidates based on the reproducibility of variant classification and the ease of assignment of new sequences. The best performance was primarily achieved with one tree-type (the strict consensus-deduplicated tree). Therefore, we focus our discussion mainly on three methods (ac.06, ac.07, and ac.08) applied to this tree-type. Phylogenetic trees for the three most common lineages (L1H, L1C, and L1A) are shown in Figure 2. RFLP and Lin+RFLP produced 82 and 142 groups, respectively, with a median of 6 and 4 sequences per group. The ac.06, ac.07, and ac.08 approaches produced 181, 151, and 115 variants, respectively, with a median of 11 to 14 sequences per variant. Only 27-30 variants were “common” variants, with at least 50 sequences. These common variants accounted for 73 – 84% of all sequences, and only 2.6 – 4.9% of sequences fell in rare variants (with fewer than 10 sequences). Bootstrap support for the ancestral node of each variant was generally high (>70%). Mean genetic distance within a variant ranged from 2.1% for ac.06 to 2.5% for ac.08, but could be as high as 4.3 to 5.3% (95^th^ percentile). In contrast, within-variant genetic distance was generally higher for RFLP (mean: 4.3%; up to 9.9% for 95^th^ percentile) and Lin+RFLP (mean: 2.5%; up to 6.6% for 95^th^ percentile). Genetic divergence from the closest-related variant was a median of 2.5 – 2.7% across the three best methods. In contrast, the median genetic divergence was only 0.5% for RFLPs and 0.7% for Lin+RFLP.

**Table 1.** Summary metrics for the average clade method with a threshold of 2 to 8% (ac.02-08) applied to deduplicated strict consensus trees (con50.dedup).

|  | RFLP | Lin+RFLP | ac.02<br>[con50.dedup] | ac.03<br>[con50.dedup] | ac.04<br>[con50.dedup] | ac.05<br>[con50.dedup] | ac.06<br>[con50.dedup] | ac.07<br>[con50.dedup] | ac.08<br>[con50.dedup] |
| --- | --- | --- | --- | --- | --- | --- | --- | --- | --- |
| Sequences per variant-median (IQR) | 6<br>(1.25-21) | 4 (1-15.75) | 4 (2-8) | 5 (2-12) | 8 (3-17) | 9 (3-20) | 11 (4-25) | 11 (4-34) | 14 (4.5-52) |
| Number variants | 82 | 142 | 815 | 428 | 284 | 219 | 181 | 151 | 115 |
| Number variants (>50) | 16 | 21 | 17 | 16 | 23 | 25 | 27 | 29 | 30 |
| Number singletons | 21 | 40 | 173 | 72 | 33 | 23 | 19 | 17 | 11 |
| % sequences in common variants | 93.3% | 87.8% | 30.4% | 48.7% | 59.9% | 68.9% | 72.9% | 79.2% | 83.6% |
| % sequences in rare variants | 2.5% | 4.3% | 34.5% | 17.1% | 10.4% | 7.4% | 4.9% | 4.0% | 2.6% |
| Bootstrap (%)-median (IQR) | 70<br>(22-76) | 74 (45-100) | 94 (67-100) | 98 (75-100) | 90 (70-100) | 98 (68-100) | 98 (70-100) | 96 (70-100) | 91 (67-100) |
| Within-variant genetic distance-mean (IQR) | 4.3%<br>(0.9-7.1%) | 2.5%<br>(0.8-3.8%) | 1.1% (0.5-1.4%) | 1.5% (0.7-1.8%) | 1.8% (1.1-2.2%) | 2.0% (1.2-2.4%) | 2.1% (1.2-2.6%) | 2.3 (1.2-3.0%) | 2.5% (1.3-3.3%) |
| Within-variant genetic distance-95<sup>th</sup> percentile | 9.90% | 6.60% | 2.80% | 3.40% | 3.90% | 4.10% | 4.30% | 4.40% | 5.30% |
| Genetic divergence from closest-related variant-median (IQR) | 0.5%<br>(0.2-1.2%) | 0.7%<br>(0.2-1.9%) | 0.8% (0.3-1.8%) | 1.7% (0.8-2.7%) | 2.0% (1.0-3.2%) | 2.2% (1.5-3.8%) | 2.5% (2.5-4.5%) | 2.5% (1.6-5.0%) | 2.7% (1.7-5.1%) |
| Assignment accuracy-internal (overall; prob.<0.2 as undetermined) | 95.3% | 93.8% | NA | NA | 99.8% | 99.4% | 99.4% | 99.2% | 99.7% |
| Assignment accuracy-external (% Nearest neighbor matching, prob.<0.2 as undetermined) | 76.5% | 80.4% | NA | NA | 95.9% | 96.6% | 96.5% | 97.6% | 96.5% |
| % farm sequence-clusters with ID change | 43.30 % | Not assessed | Not assessed | Not assessed | Not assessed | Not assessed | 8.70% | 8.70% | 6.50% |

**Figure 2.**
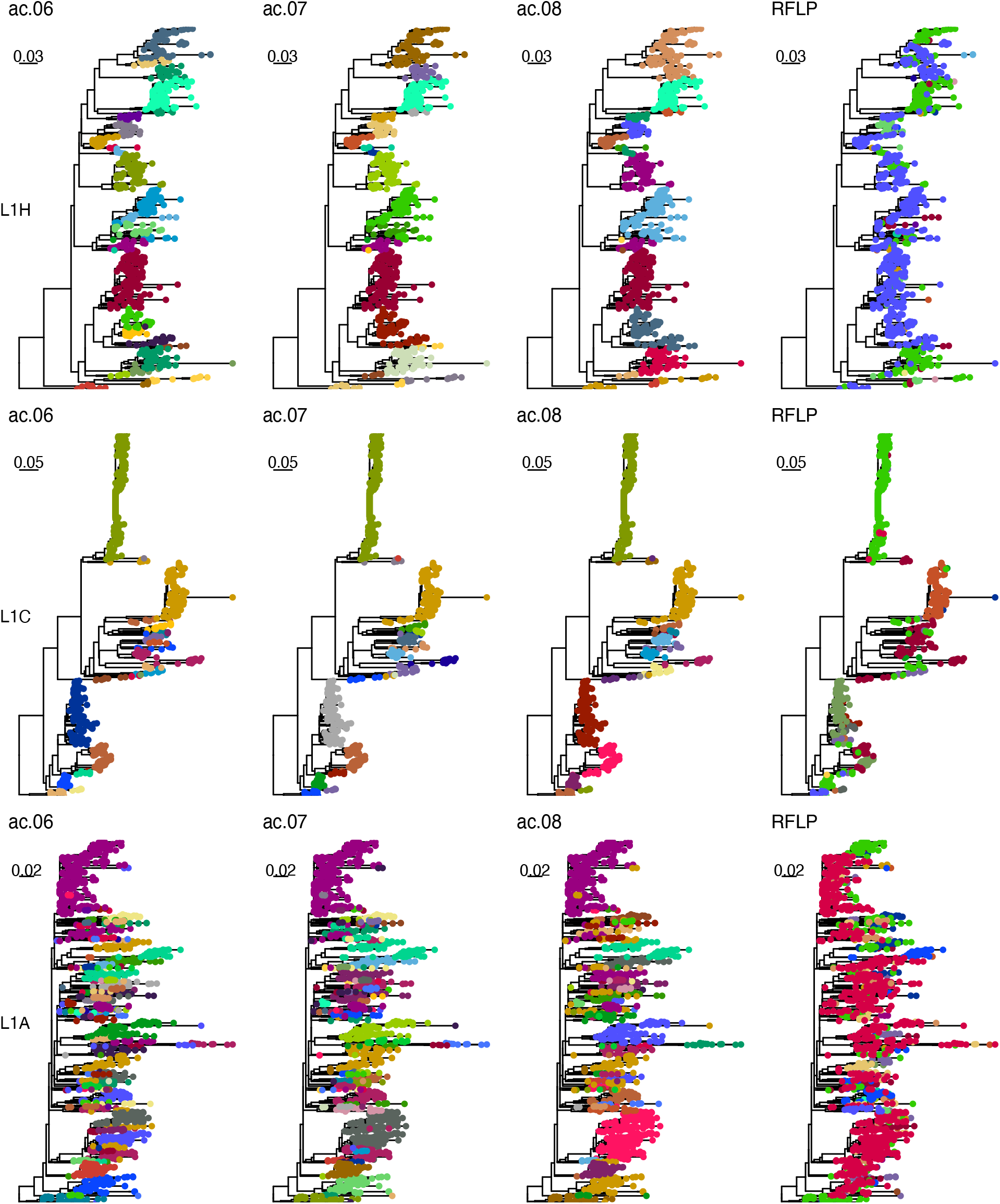
Example trees. Phylogenetic trees for sub-lineage L1H (top row), L1C (middle row), and L1A (bottom row). Colors in the first, second, third, and fourth columns represent classifications with the ac.06, ac.07, ac.08, and RFLP methods. Sequences with the same RFLP-type are denoted with the same color across all three lineages.

### 3.2 Reproducibility of classification amongst sets of data

We performed several analyses to determine the extent to which the variant classification produced above could be replicated with different sets of data. When variant clustering was performed on duplicate phylogenetic trees from different IQ-TREE runs or on a detailed sub-lineage L1C tree with 15 years of data, concordance between the classifications produced for each tree was quantified through the Jaccard index (Figure 3a). Index values of >0.85 are generally considered highly stable (41). The Jaccard index ranged between 0.78 and 0.97 for duplicate trees (black points in Figure 3a), and between 0.31 and 0.95 for the L1C trees (red in Figure 3a). For the latter, the Jaccard indices improved to 0.83 and 0.95 when considering trees with ≥6% threshold and were notably poor for lower thresholds, indicating a lack of reproducibility when the threshold was set too low.

In another reproducibility analysis, trees were constructed with 10 random subsets of the short-term dataset, and then the variant IDs from the full analysis were annotated onto the trees. We quantified the proportion of sequences with a matching nearest neighbor as well as clade purity across each of the ten trees. All methods achieved high nearest neighbor matching, with >94% of sequences having a nearest neighbor that had a matching variant ID (black in Figure 3b). Only ∼80% of sequences had a matching nearest neighbor when using RFLP or Lineage+RFLP. Median clade purity ranged from 79-94% across approaches, with higher purities of nearly 90% or more achieved for lower thresholds (4 and 5%) and for approaches utilizing the strict consensus method on de-duplicated data (red in Figure 3b). Median clade purities were 49% and 69% for RFLP and Lineage+RFLP, respectively.

All approaches were able to capture the *a priori* defined outbreak variant known as L1C-1-4-4, with Jaccard indices of 0.96 – 0.98. RFLP and Lineage+RFLP achieved only a concordance of 0.28 and 0.76, respectively, indicating that these labels did not reliably capture sequences associated with this outbreak.

Although variants were defined on trees built via IQ-TREE, Bayesian methods such as BEAST are often considered the most robust approach. Therefore, we assessed whether the variants produced on the IQ-TREEs also formed clusters with high purity on Bayesian trees. The median clade purity for variants on time-scaled Bayesian trees was essentially 1 in all cases, but the lower bound of the interquartile range was more variable (black in Figure 3c) and was particularly low for RFLP and Lineage+RFLP. Median clade purity was also more variable when considering only Lineage 1A (red in Figure 3c), which was consistently the most problematic lineage in all analyses.

**Figure 3.**
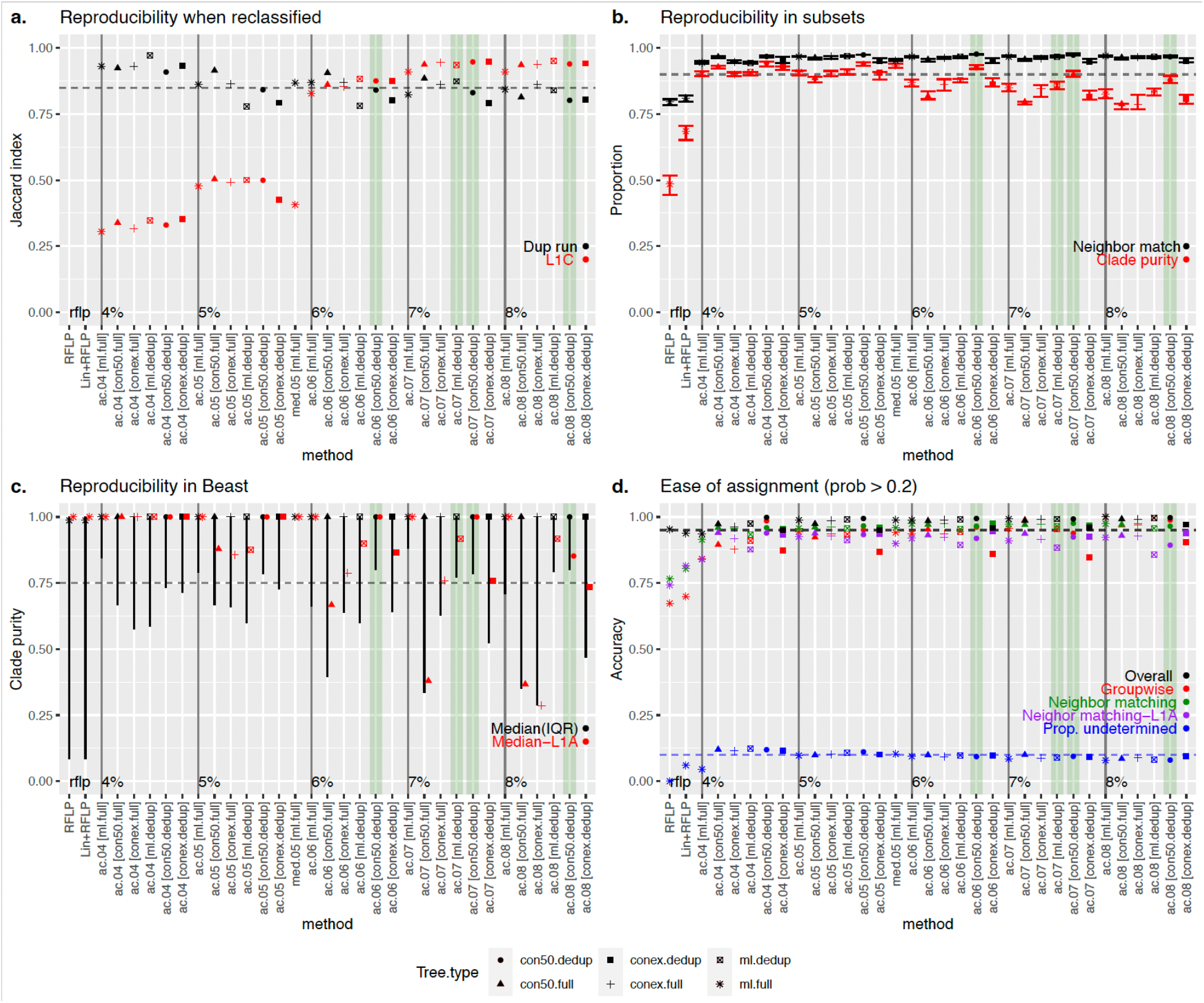
Comparison of all approaches that produced a median variant size >5 sequences/variant. Best performing methods are highlighted in green. a) Reproducibility when reclassified: Concordance (as measured by the Jaccard index) between the classifications produced on duplicate IQ-TREE runs performed with the same data (black) or between the full analysis and the more detailed set of L1C sequences (red). b) Reproducibility in 10 random subsets of data: Neighbor match (black)-Proportion of sequences whose nearest neighbor in the tree had a matching variant ID (black), and average proportion of sequences in a clade that shared the same ID (clade-purity, red). c) Reproducibility in BEAST: Median and interquartile range of the clade purity of all variants (black) or L1A variants (red) in Bayesian phylogenies constructed using BEAST. d) Ease of assignment: Overall (black) and group-wise accuracy (red) of the assignment algorithm’s ability to correctly assign variant IDs to new sequences (internal test set). Proportion of newly assigned sequences (external test set) whose nearest neighbor in the tree had a matching variant ID (green-all sequences; purple-L1A sequences). Proportion of sequences whose variant ID was undetermined (i.e., confidence in assignment was >0.2 probability). In all plots, the dotted line represents the desired value for each assessment. Tree type is represented by shape.

### 3.3 Ease of classification

For each of the 33 classification approach considered, we trained a random forest algorithm to assign variant IDs to sequences. Model performance was evaluated with an internal test set (most recent 10% of data from the short-term dataset) and an external test set (sequences that were not included in the original analyses, Figure 4). Predictions from the trained random forest algorithms include the probabilities of the first, second, and third most likely variant IDs for a given sequence, with the highest probability ID being assigned to the sequence for downstream analyses of predictive performance. However, in some cases, the highest probability ID was quite low, indicating that the model had poor confidence in the assignment. Therefore, we tested two thresholds (prob < 0.2 or prob <0.6) for calling sequences “undetermined” rather than assigning them to an ID. The 0.2 threshold was determined by taking the median probability for mis-classified sequences in the internal test set (ac.06). We then assessed the improvement in overall and group-wise accuracy in the internal test data when low-certainty sequences were removed from the calculation. For the external test data, we evaluated improvements in nearest neighbor matching overall and for Lineage 1A (the lineage with poorest predictive accuracy). We also tabulated the percentage of sequences that were undetermined (Figure 3d).

Both uncertainty thresholds resulted in marked improvement in predictive performance across the board, except for the ac.04 methods and trees built with the extended consensus method (de-duplicated) which never achieved comparable accuracies as other approaches. The 0.2 threshold improved the internal test set overall accuracy to 97.4 – 100% and groupwise accuracy to 92.3 - 100%, and external test set nearest neighbor matching to 95.4 – 97.6% (Supplementary Figure S1). The 0.6 threshold improved the internal test set overall accuracy 99.2-100% and groupwise accuracy by 97.3 - 100%, and external test set nearest neighbor matching to 98.8-99.7% (Supplementary Figure S1). While the 0.6 probability threshold resulted in slightly higher accuracies, it also resulted in a high percentage (25%) of undetermined sequences in the external test set. With comparable predictive performance, the 0.2 threshold resulted in just 10% of sequences classified as undetermined (Figure 3d). Therefore, a probability threshold of <0.2 for calling sequences undetermined was applied to predictions made by the random forest algorithm. Even for the problematic sub-lineage L1A, the top 10 approaches all achieved >93% nearest neighbor matching in the external test set (Supplementary Figure 1-2). Recall and precision ranged from 99.6% - 100% and 99.9 – 100%, respectively.

**Figure 4.**
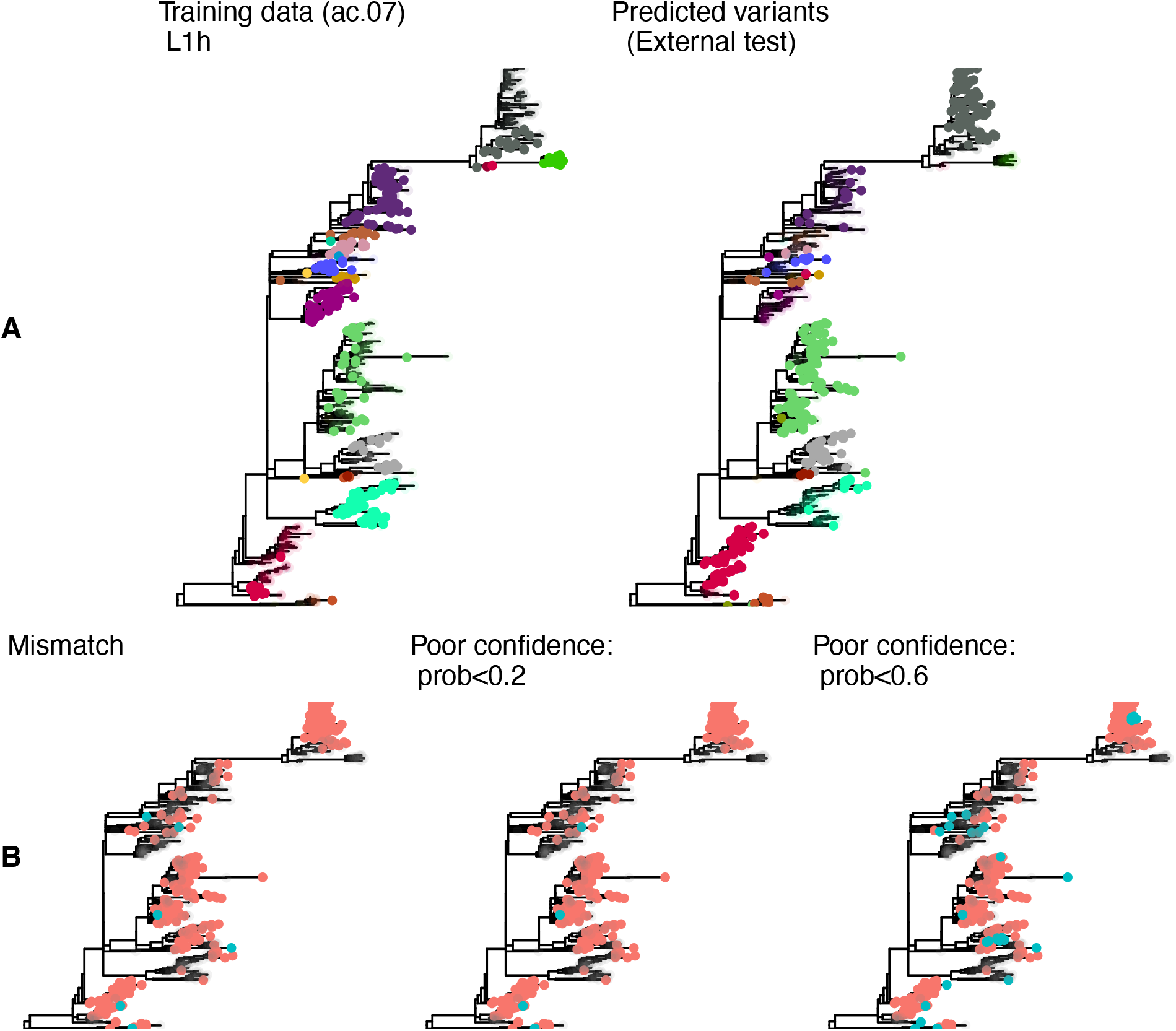
Phylogenetic trees for sub-lineage L1H constructed with sequences from the UMN Veterinary Diagnostic Laboratory. A) colors represent the variant ID for sequences present in the training data, with uncolored tips indicating test set sequences (left); and the predicted ID for the external test set, with uncolored tips representing training sequences (right). The predicted variant ID was considered correct if the nearest training sequence had the same variant ID. B) Mismatches (blue tips) are showin in the left tree, and predictions flagged as poor confidence (blue tips) are shown for the 0.2 threshold (middle tree) and 0.6 threshold (right tree).

### 3.4 Selection of best classification approaches

For the reproducibility analysis, the best performing approaches were defined as those that achieved a Jaccard index of >0.85 for both re-classification on the duplicate run and on the extended L1C analysis (*criteria 1)*. Nine of 33 approaches met the criteria. For the subset analysis, the best performing approaches were defined as those that achieved both clade purity and nearest neighbor matching of >0.90 (*criteria 2*). 14 approaches met this criterion. For the reproducibility analysis using trees generated by BEAST, the best performing approaches were defined as those in which the lower bound of the interquartile range for clade purity was >0.75 (*criteria 3*). 10 approaches met this criterion. Finally, for ease of classification, we defined the best performing approaches as those with >0.95 overall accuracy in the internal test data as well as >0.90 for both the mean groupwise accuracy in the internal test set and nearest neighbor matching in the external test set (*criteria 4*). 21 approaches met this criterion. Values that missed the threshold by <0.01 were allowed for all criteria. Only one approach (variant.06.ac.dedup.con) met all criteria, which was the approach with a 6% threshold for average clade method applied to a strict consensus tree with the deduplicated dataset. An additional three approaches satisfied three of the four criteria and missed only one criterion by no more than 0.05 (Figure 3). Given that one tree type (strict consensus tree with de-duplicated sequences) accounted for three of four of the best approaches, we proceeded with that tree type for the remaining analyses.

### 3.5 Farm-level occurrence of variants

We used the long-term dataset (which spans approximately 11 years of sequences) to tabulate the number of MSHMP-participating farms, production systems, and U.S. states in which each variant was detected. These summaries excluded rare variants (<10 sequences), which accounted for 3.5, 2.4, and 1.8% of sequences, respectively for ac.06, ac.07, and ac.08. Variants (ac.06, ac.07, and ac.08) were found in a median of 8 to 9 farms (max 24), 3 (max 6) production systems, and 2 (max 3) states.

To assess the stability of variant classification during micro-evolution that may occur during virus circulation on a farm, 73 farms with at least 4 sequences in a single year were identified from the long-term dataset. From these, 587 sequences were available (4 – 43 sequences per farm, with some farms meeting the yearly criteria multiple times). For each pair of variants that occurred on the same farm, we measured the maximum genetic distance and maximum divergence time between those sequences to help identify situations in which sequences from a single sequence-cluster (characterized by low genetic distance and short divergence times) on a farm were classified as two distinct IDs (Supplementary Figure S3). For ac.06 and ac.07, all variant pairs with maximum genetic distance <0.05 and/or divergence time <2 years were manually inspected on the trees as to whether those sequences would be more accurately represented by a single variant ID (i.e., they cluster together in the tree, Figure 1). Based on this manual inspection, a merging threshold of <0.02 genetic distance and <3 years divergence time was applied to all pairs of variants from the same farm. Variant pairs from the same farm that met both these conditions were flagged as farm sequence-clusters in which an ID change occurred as a result of micro-evolution (though we cannot rule-out the possibility of two separate introductions of closely related viruses onto the farm). An ideal classification system should minimize the occurrence of such ID changes. The percent of farm sequence-clusters with an ID change was 8.7%, 8.7%, and 6.5% for ac.06, ac.07, and ac.08, respectively. In contrast, ∼43% of farm sequence-clusters had an RFLP change.

## 4. Discussion

Due to the rapid diversification and spread of PRRSV-2, current classification systems based on lineages/sub-lineage systems do not provide adequate resolution on PRRSV-2 diversity, and RFLP-typing does not reliably group together related sequences. In this paper, we compare the performance of different phylogenetic clustering approaches on a large dataset of PRRSV-2 sequences from the United States. One of our key objectives was to evaluate alternatives for fine-scale classification of PRRSV-2. We first identified approaches for delineating variants within phylogenetic trees that were statistically supported and provide robust and reproducible results when analyses were repeated with various subsets of data. We found that our best-performing approaches achieved high consistency in which sequences were identified as belonging to the same genetic variant across multiple analyses, thus demonstrating the robustness of these approaches.

In addition, we show a substantial advantage of the new approaches over RFLPs and Lin+RFLPs in grouping together highly related sequences and disaggregating genetically more distant sequences. This demonstrates that these alternative systems address the shortcomings of RFLP-based classification, where genetically similar sequences often receive different RFLP-types and genetically distant sequences have the same RFLP-type. Likewise, these new approaches provide a more granular sequence classification that lineages and sub-lineages, thus can be used to quickly indicate an emergent variant clade. In addition, we also found that our best-performing approaches were able to minimize ID changes that occur during a single PRRSV-2 circulation event on a farm, with just 6-9% of circulation events having an ID change as compared to >40% of on-farm circulation events having an RFLP-type change.

Our comparison of clustering methods lays the foundation for fine-scale classification of PRRSV-2 that meets the needs of animal professionals utilizing sequence data as part of disease monitoring and management. That being said, any nomenclature based on ORF5 sequences will not fully represent the evolutionary ancestry or phenotypic expression of a given virus, as recombination across the genome may alter the evolutionary relationships between different parts of the genome. In some cases, recombinant clades (i.e., groups of sequences likely descended from a recombinant ancestor) appear as divergent groups in ORF5 phylogenies, even if the recombination event occurred outside of the ORF5 gene (36, 42). Hence, their unique evolutionary trajectory is sometimes discernable in ORF5 phylogenies in instances where the recombination event produced numerous descendants in the viral population, but this is not always true (43). Whole genome sequencing would be required to fully characterize these recombination events, but a fine-scale classification approach based on ORF5 would be able to discern these distinct groups in many cases (42).

It is unknown if genetic diversity captured by the variants identified here translate into phenotypic diversity of the virus, either at an antigenic or virulence level. While the ORF5 gene is immunologically important, other parts of the genome contribute to the antigenicity and virulence (20, 44, 45). Whole genome data would be needed to understand the interplay between genotype and phenotype, and observed clinical manifestations are also influenced by external factors (e.g., co-infections); science has not yet progressed to the point that we can predict phenotype from whole genomes for PRRSV (22). That being said, sequencing conducted by animal health professionals is often conducted for epidemiological monitoring purposes, and this informed the level of granularity that we tried to achieve in this analysis.

While we utilized a data set of PRRV-2 sequences that is representative of ∼70% of the U.S. sow population, with even representation across all major pig producing regions (10), it is possible that there are pockets of genetic diversity not represented in our dataset that may constitute distinct variants. A more comprehensive dataset could mitigate this possibility. In addition, our analysis is U.S. specific, and not representative of global PRRSV-2 diversity. Due to the granularity of the variants defined here (typically <2.5% within-variant genetic distances), PRRSV-2 in other countries would likely be sufficiently diverged to be classified as distinct variants from those of the U.S., but the ability to define PRRSV-2 variants in each country is dependent on sequence availability.

Finally, the farm-level analysis was dependent on available retrospective data from routine veterinary care, and therefore sequences assigned to farm-specific sequence-clusters (i.e., closely related sequences from a single farm that likely represent a single PRRSV circulation event) were not collected in a systematic manner. Indeed, farms that met our criteria for sequence availability may favor the inclusion of farms experiencing atypical circulation events (i.e., those with prolonged circulation or more severe clinical signs) due to biases in what is selected by veterinarians for sequencing. That may contribute to the surprisingly high percentage of farm sequence-clusters that had RFLP-type changes, which may be an overestimate as a result of these biases. That being said, ID changes pose a particular challenge in farms with prolonged circulation vents, due to greater elapsed time for micro-evolution on the farm.

More generally, our comparison of different methodologies for clustering sequences within phylogenies highlights several insights that may be applicable to other RNA viruses. Phylogenies built with many closely related sequences, especially when a relatively short marker gene is used, can result in trees with low bootstrap support and unstable topologies (27). Here, we show that improved reproducibility of the clustering analysis can be achieved if a strict consensus tree is used, as nodes with low support are collapsed; only nodes with high bootstrap support are retained in the tree. We also show that the average clade method consistently outperforms the other methods tested.

Through identifying methodology that group together related sequences in a robust and reproductible manner, this work lays the foundation for fine-scale classification of PRRSV-2 in the U.S. Next steps that build upon this work are to test the performance and robustness of this nomenclature when performed on a rolling basis, which will evaluate the ability of a new classification system to accommodate expanding genetic diversity as the virus continues to evolve. Additional next steps include defining a standardized system to label genetic variants, developing procedures for prospective implementation, and establishing mechanisms for defining variants in international contexts. In these future steps, input from stakeholders is crucial to establish a system that meets the needs of diagnostic labs and animal health professionals.

## 6. Author Contributions

KV designed the research, performed the analysis, and wrote the paper. NP, MK, and IP performed some analyses, contributed to interpretation, and edited the paper. DL, GT, JZ, TA, MZ, CC and DH contributed to research design, results interpretation, and manuscript editing.

## Acknowledgments

The authors would like to thank members of the AASV PRRSV nomenclature working group, including Andreia Arruda, Srijita Chandra, Eric Nelson, Tom Petznick, Melanie Prarat, Mark Schwartz, Jessica Seate, Gustavo Silva, Joel Sparks, and Paul Yeske, The authors would like to thank the Morrison Swine Health Monitoring Project (MSHMP) participants and the Veterinary Diagnostic Laboratory, University of Minnesota, for sharing its PRRSV-2 genetic sequences. Mention of trade names or commercial products in this article is solely for the purpose of providing specific information and does not imply recommendation or endorsement by the U.S. Department of Agriculture (USDA). The funders had no role in study design, data collection and interpretation, or the decision to submit the work for publication. The findings and conclusions in this publication are those of the authors and should not be construed to represent any official USDA or U.S. Government determination or policy. USDA is an equal opportunity provider and employer.

## Supplementary material

**Supplementary figure S1.**
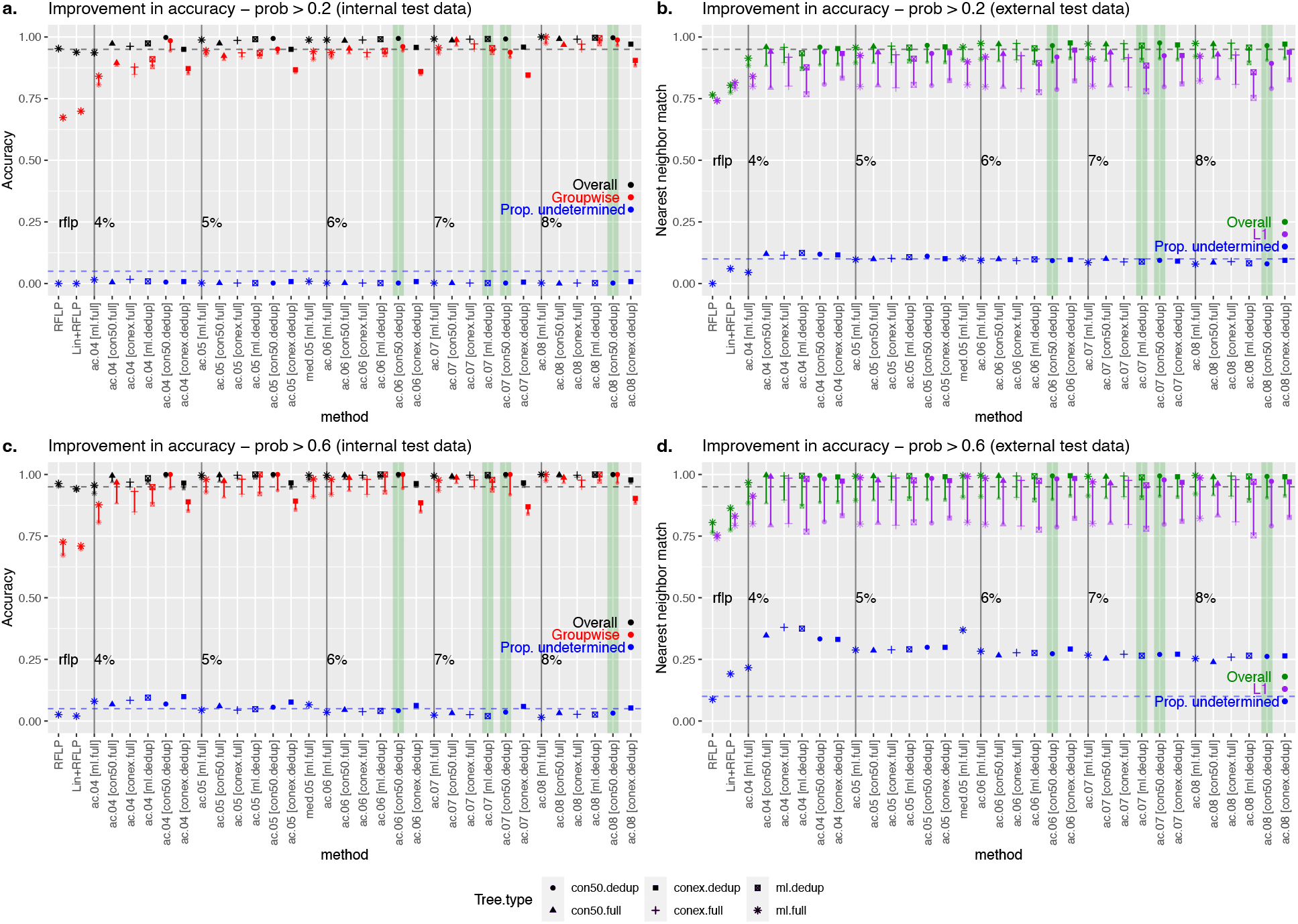
Assignment accuracy improvements when sequences are classified as “undetermined” if their assignment probability is <0.2 (panels a and b) or 0.6 (panels c and d). Best performing methods are highlighted in green. (a and c): Improvements in the overall (black) and groupwise accuracy (red), and the proportion of sequences that are undetermined (blue) in the internal test data. (b and d): Improvements in accuracy for the external test data, as measured by nearest neighbor matching: overall (green), lineage 1A (purple), and proportion of sequences that are undetermined (blue). In all plots, the dotted line represents the desired value for each assessment. Tree type is represented by shape.

**Supplementary figure S2.**
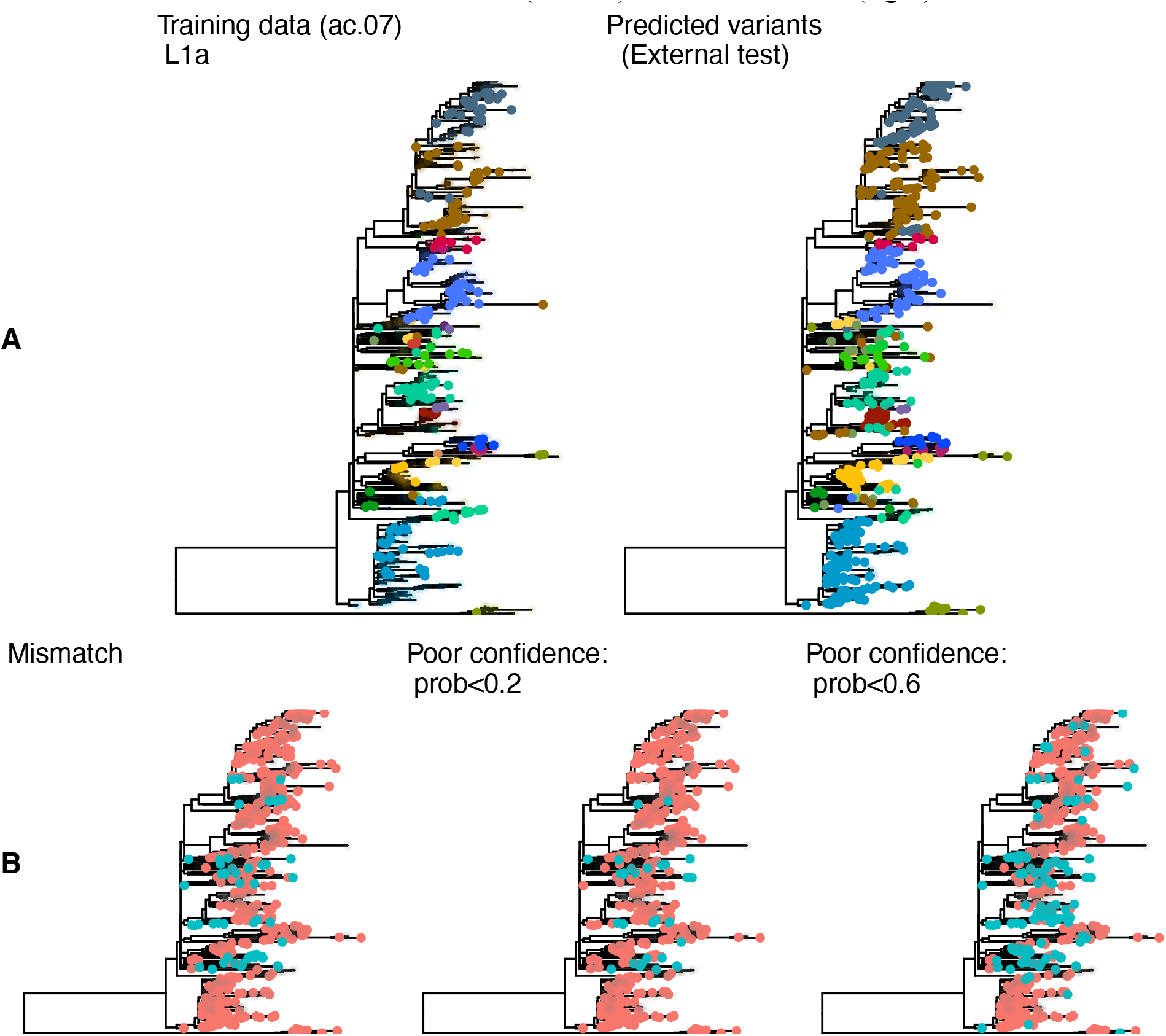
Phylogenetic trees for sub-lineage L1A constructed with sequences from the UMN Veterinary Diagnostic Laboratory. A. colors represent the variant ID for sequences present in the training data, with transparaent colors indicating test set sequences (left); and the predicted ID for the external test set, with transparent colors representing training sequences (right). The predicted ID was considered correct if the nearest training sequence was the same ID. B. Mismatches are showin in the left tree, and predictions flagged as poor confidence are shown for the 0.2 threshold (middle) and 0.6 threshold (right)

**Supplementary figure S3.**
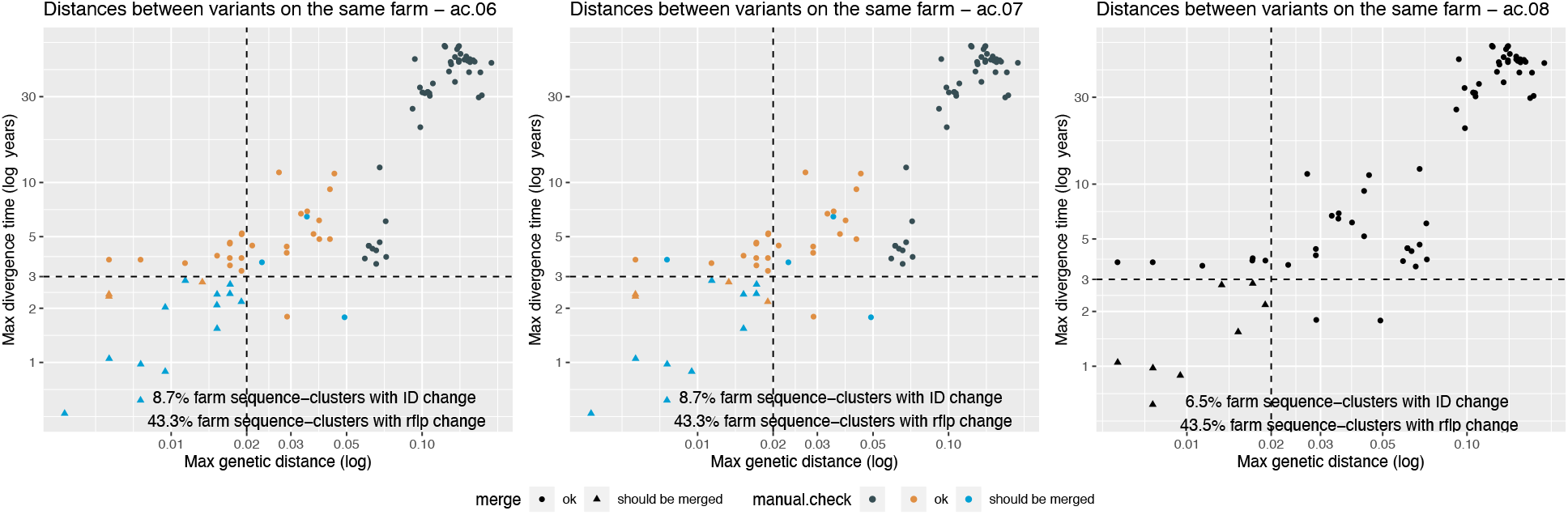
Scatterplot of the maximum genetic (patristic) distance and divergence time for each pair of variants detected on the same farm for the ac.06, ac.07, and ac.08 methods. Colors indicate whether the farm sequence-cluster was manually checked as to whether the two sequences belonging to the two IDs should be merged (they form a single cluster; blue) or whether they are ok represented as two clusters (orange). Those with genetic distance >0.05 were not manually checked, as it was assumed that the two variant IDs were not closely related. Hashed lines indicate thresholds used to discriminate between clusters that should or should not be merged. Shape indicates those that should (triangle) or should not (circle) be merged based solely on threshold (without manual checks).

