## Supplementary Table S1 for "Phylogenetic-based methods for fine-scale classification of PRRSV-2 ORF5 sequences: a comparison of their robustness and reproducibility"

| | Approach | Tree type / full vs. deduplicated data | Threshold | Clustering method | In best 33 candidates | Sequences per variant (median) | Sequences per variant (IQR-25%) | Sequences per variant (IQR-75%) | Number variants | Number variants (>50) | Number singletons | % sequences in common variants | % sequences in rare variants | Bootstrap (%)-median | Within-variant genetic distance-mean | Within-variant genetic distance-95th percentile | Genetic divergence from closest-related variant-median | accuracy-internal (overall; prob.<0.2 as undetermined) | Assignment accuracy-internal (group wise, prob.<0.2 as undetermined) | accuracy-external (% Nearest neighbor matching, prob.<0.2 as undetermined) | % farm sequence-clusters with ID change | Jaccard index (duplicate run) | Jaccard index (L1C) | Nearest neighbor matching in subsets (median) | Nearest neighbor matching in subsets(IQR-25%) | Nearest neighbor matching in subsets (IQR-75%) | Clade purity in subsets (median) | Clade purity in subsets (IQR-25%) | Clade purity in subsets (IQR-75%) | Clade purity with BEAST (median) | Clade purity with BEAST (IQR-25%) | Clade purity with BEAST (IQR-75%) |
|---|---|---|---|---|---|---|---|---|---|---|---|---|---|---|---|---|---|---|---|---|---|---|---|---|---|---|---|---|---|---|---|---|
| 1 | rflp | ml.full | | rflp | yes | 6 | 1.25 | 21 | 82 | 16 | 21 | 0.933 | 0.025 | 52 | 0.043 | 0.099 | 0.01 | 0.953 | 0.673 | 0.765 | 43.3 | #N/A | #N/A | 0.796 | 0.784 | 0.806 | 0.485 | 0.444 | 0.517 | 0.987 | 0.085 | 1 |
| 2 | rflp.lin | ml.full | | rflp-lin | yes | 4 | 1 | 15.75 | 142 | 21 | 40 | 0.878 | 0.043 | 81 | 0.025 | 0.066 | 0.016 | 0.938 | 0.699 | 0.804 | #N/A | #N/A | #N/A | 0.811 | 0.796 | 0.821 | 0.685 | 0.652 | 0.705 | 0.987 | 0.085 | 1 |
| 3 | ac.05 | ml.full | 05 | ac | yes | 9 | 3 | 22.75 | 202 | 28 | 24 | 0.718 | 0.064 | 96 | 0.019 | 0.036 | 0.029 | 0.988 | 0.944 | 0.957 | #N/A | 0.862 | 0.477 | 0.967 | 0.966 | 0.97 | 0.909 | 0.897 | 0.914 | 1 | 0.789 | 1 |
| 4 | ac.04 | ml.full | 04 | ac | yes | 7 | 2 | 18 | 304 | 24 | 47 | 0.569 | 0.105 | 95.5 | 0.016 | 0.034 | 0.023 | 0.936 | 0.841 | 0.914 | #N/A | 0.931 | 0.305 | 0.945 | 0.942 | 0.951 | 0.9 | 0.894 | 0.911 | 1 | 0.845 | 1 |
| 5 | med.05 | ml.full | 05 | med | yes | 7 | 2 | 17.5 | 263 | 21 | 0 | 0.625 | 0.101 | 100 | 0.012 | 0.022 | 0.025 | 0.988 | 0.941 | 0.959 | #N/A | 0.868 | 0.407 | 0.952 | 0.95 | 0.96 | 0.933 | 0.924 | 0.94 | 1 | 1 | 1 |
| 6 | ac.06 | ml.full | 06 | ac | yes | 13 | 4 | 41 | 153 | 30 | 15 | 0.778 | 0.042 | 95 | 0.02 | 0.04 | 0.033 | 0.988 | 0.937 | 0.974 | #N/A | 0.869 | 0.828 | 0.967 | 0.965 | 0.97 | 0.869 | 0.854 | 0.88 | 1 | 0.662 | 1 |
| 7 | ac.07 | ml.full | 07 | ac | yes | 14 | 4 | 55 | 119 | 32 | 12 | 0.839 | 0.031 | 93 | 0.021 | 0.043 | 0.038 | 0.992 | 0.956 | 0.972 | #N/A | 0.824 | 0.910 | 0.967 | 0.966 | 0.972 | 0.849 | 0.836 | 0.864 | 1 | 0.881 | 1 |
| 8 | ac.08 | ml.full | 08 | ac | yes | 16 | 5 | 66 | 97 | 29 | 10 | 0.867 | 0.024 | 91 | 0.023 | 0.043 | 0.042 | 1 | 1 | 0.974 | #N/A | 0.843 | 0.910 | 0.97 | 0.968 | 0.973 | 0.825 | 0.81 | 0.843 | 1 | 0.708 | 1 |
| 9 | sl.05 | ml.full | 05 | sl | no | 2 | 1 | 6.25 | 320 | 20 | 140 | 0.779 | 0.102 | 100 | 0.01 | 0.021 | 0.033 | | #N/A | #N/A | #N/A | #N/A | #N/A | 0.972 | 0.964 | 0.976 | 0.975 | 0.967 | 0.978 | #N/A | #N/A | #N/A |
| 10 | len.03 | ml.full | 03 | len | no | 2 | 1 | 6 | 212 | 13 | 101 | 0.874 | 0.069 | 100 | 0.009 | 0.021 | 0.032 | #N/A | #N/A | #N/A | #N/A | #N/A | #N/A | 0.955 | 0.95 | 0.958 | 0.97 | 0.965 | 0.974 | #N/A | #N/A | #N/A |
| 11 | len.04 | ml.full | 04 | len | no | 2 | 1 | 6 | 212 | 13 | 101 | 0.874 | 0.069 | 100 | 0.01 | 0.027 | 0.038 | #N/A | #N/A | #N/A | #N/A | #N/A | #N/A | 0.98 | 0.973 | 0.983 | 0.985 | 0.978 | 0.986 | #N/A | #N/A | #N/A |
| 12 | len.05 | ml.full | 05 | len | no | 2 | 1 | 8.5 | 143 | 13 | 61 | 0.899 | 0.05 | 100 | 0.012 | 0.037 | 0.045 | #N/A | #N/A | #N/A | #N/A | #N/A | #N/A | 0.984 | 0.981 | 0.988 | 0.973 | 0.964 | 0.975 | #N/A | #N/A | #N/A |
| 13 | ac.03 | ml.full | 03 | ac | no | 5 | 2 | 12 | 482 | 17 | 104 | 0.444 | 0.185 | 95 | 0.014 | 0.031 | 0.018 | #N/A | #N/A | #N/A | #N/A | #N/A | 0.156 | 0.913 | 0.901 | 0.92 | 0.931 | 0.926 | 0.947 | #N/A | #N/A | #N/A |
| 14 | med.03 | ml.full | 03 | med | no | 2 | 1 | 5 | 825 | 16 | 350 | 0.439 | 0.259 | 100 | 0.007 | 0.014 | 0.016 | #N/A | #N/A | #N/A | #N/A | #N/A | 0.141 | 0.919 | 0.909 | 0.928 | 0.954 | 0.945 | 0.961 | #N/A | #N/A | #N/A |
| 15 | sl.03 | ml.full | 03 | sl | no | 2 | 1 | 4 | 679 | 17 | 336 | 0.591 | 0.21 | 100 | 0.006 | 0.015 | 0.022 | #N/A | #N/A | #N/A | #N/A | #N/A | #N/A | 0.946 | 0.944 | 0.952 | 0.97 | 0.967 | 0.975 | #N/A | #N/A | #N/A |
| 16 | med.04 | ml.full | 04 | med | no | 2 | 1 | 9 | 558 | 17 | 192 | 0.488 | 0.181 | 100 | 0.009 | 0.018 | 0.02 | #N/A | #N/A | #N/A | #N/A | #N/A | 0.321 | 0.956 | 0.952 | 0.959 | 0.937 | 0.933 | 0.947 | #N/A | #N/A | #N/A |
| 17 | sl.04 | ml.full | 04 | sl | no | 2 | 1 | 6 | 462 | 22 | 199 | 0.688 | 0.151 | 100 | 0.008 | 0.017 | 0.028 | #N/A | #N/A | #N/A | #N/A | #N/A | #N/A | 0.962 | 0.958 | 0.964 | 0.971 | 0.967 | 0.973 | #N/A | #N/A | #N/A |
| 18 | sl.02 | ml.full | 02 | sl | no | 1 | 1 | 3 | 1100 | 14 | 601 | 0.465 | 0.32 | 100 | 0.004 | 0.01 | 0.017 | #N/A | #N/A | #N/A | #N/A | #N/A | #N/A | 0.915 | 0.902 | 0.921 | 0.968 | 0.968 | 0.971 | #N/A | #N/A | #N/A |
| 19 | len.02 | ml.full | 02 | len | no | 1 | 1 | 3 | 677 | 16 | 379 | 0.659 | 0.19 | 100 | 0.006 | 0.016 | 0.021 | #N/A | #N/A | #N/A | #N/A | #N/A | #N/A | 0.92 | 0.917 | 0.927 | 0.962 | 0.96 | 0.966 | #N/A | #N/A | #N/A |
| 20 | ac.02 | ml.full | 02 | ac | no | 3 | 1 | 8 | 860 | 10 | 233 | 0.268 | 0.355 | 93.5 | 0.011 | 0.027 | 0.012 | #N/A | #N/A | #N/A | #N/A | #N/A | #N/A | 0.856 | 0.845 | 0.865 | 0.941 | 0.937 | 0.949 | #N/A | #N/A | #N/A |
| 21 | lenc.05 | ml.full | 05 | lenc | no | 3 | 1 | 9 | 491 | 30 | 185 | 0.585 | 0.159 | 100 | 0.01 | 0.029 | 0.02 | #N/A | #N/A | #N/A | #N/A | #N/A | #N/A | 0.895 | 0.892 | 0.901 | 0.917 | 0.909 | 0.927 | #N/A | #N/A | #N/A |
| 22 | lenc.03 | ml.full | 03 | lenc | no | 2 | 1 | 5 | 1061 | 18 | 483 | 0.311 | 0.348 | 100 | 0.006 | 0.017 | 0.015 | #N/A | #N/A | #N/A | #N/A | #N/A | #N/A | 0.855 | 0.853 | 0.868 | 0.94 | 0.932 | 0.944 | #N/A | #N/A | #N/A |
| 23 | lenc.04 | ml.full | 04 | lenc | no | 2 | 1 | 7 | 731 | 29 | 300 | 0.444 | 0.238 | 99 | 0.008 | 0.023 | 0.017 | #N/A | #N/A | #N/A | #N/A | #N/A | #N/A | 0.867 | 0.862 | 0.873 | 0.917 | 0.906 | 0.927 | #N/A | #N/A | #N/A |
| 24 | lenc.02 | ml.full | 02 | lenc | no | 1 | 1 | 3 | 1753 | 11 | 900 | 0.128 | 0.548 | 100 | 0.004 | 0.012 | 0.011 | #N/A | #N/A | #N/A | #N/A | #N/A | #N/A | 0.843 | 0.831 | 0.849 | 0.96 | 0.952 | 0.962 | #N/A | #N/A | #N/A |
| 25 | med.02 | ml.full | 02 | med | no | 1 | 1 | 3 | 1507 | 9 | 785 | 0.266 | 0.479 | 100 | 0.004 | 0.009 | 0.012 | #N/A | #N/A | #N/A | #N/A | #N/A | #N/A | 0.855 | 0.848 | 0.889 | 0.969 | 0.965 | 0.974 | #N/A | #N/A | #N/A |
| 26 | 02.ac | conex.full | 02 | ac | no | 3 | 1 | 8 | 845 | 12 | 217 | 0.288 | 0.348 | 93 | 0.01 | 0.027 | 0.014 | #N/A | #N/A | #N/A | #N/A | #N/A | #N/A | 0.858 | 0.843 | 0.868 | 0.949 | 0.945 | 0.953 | #N/A | #N/A | #N/A |
| 27 | 02.len | conex.full | 02 | len | no | 1 | 1 | 3 | 674 | 16 | 377 | 0.656 | 0.189 | 100 | 0.006 | 0.017 | 0.022 | #N/A | #N/A | #N/A | #N/A | #N/A | #N/A | 0.922 | 0.912 | 0.927 | 0.962 | 0.96 | 0.964 | #N/A | #N/A | #N/A |
| 28 | 02.lenc | conex.full | 02 | lenc | no | 2 | 1 | 3 | 1702 | 11 | 847 | 0.133 | 0.54 | 99 | 0.004 | 0.012 | 0.012 | #N/A | #N/A | #N/A | #N/A | #N/A | #N/A | 0.859 | 0.847 | 0.861 | 0.961 | 0.959 | 0.964 | #N/A | #N/A | #N/A |
| 29 | 02.med | conex.full | 02 | med | no | 1 | 1 | 3 | 1447 | 8 | 727 | 0.27 | 0.47 | 100 | 0.004 | 0.009 | 0.012 | #N/A | #N/A | #N/A | #N/A | #N/A | #N/A | 0.858 | 0.852 | 0.882 | 0.97 | 0.969 | 0.976 | #N/A | #N/A | #N/A |
| 30 | 02.sl | conex.full | 02 | sl | no | 1 | 1 | 3 | 1096 | 13 | 602 | 0.468 | 0.315 | 100 | 0.004 | 0.01 | 0.017 | #N/A | #N/A | #N/A | #N/A | #N/A | #N/A | 0.918 | 0.905 | 0.923 | 0.97 | 0.969 | 0.971 | #N/A | #N/A | #N/A |
| 31 | 03.ac | conex.full | 03 | ac | no | 5 | 2 | 12.75 | 478 | 18 | 94 | 0.449 | 0.177 | 92 | 0.014 | 0.033 | 0.019 | #N/A | #N/A | #N/A | #N/A | #N/A | 0.156 | 0.903 | 0.892 | 0.907 | 0.938 | 0.925 | 0.941 | #N/A | #N/A | #N/A |
| 32 | 03.len | conex.full | 03 | len | no | 1 | 1 | 4 | 379 | 15 | 197 | 0.795 | 0.117 | 100 | 0.008 | 0.021 | 0.03 | #N/A | #N/A | #N/A | #N/A | #N/A | #N/A | 0.957 | 0.953 | 0.96 | 0.971 | 0.966 | 0.974 | #N/A | #N/A | #N/A |
| 33 | 03.lenc | conex.full | 03 | lenc | no | 2 | 1 | 5 | 1081 | 16 | 482 | 0.301 | 0.358 | 100 | 0.006 | 0.017 | 0.015 | #N/A | #N/A | #N/A | #N/A | #N/A | #N/A | 0.876 | 0.867 | 0.877 | 0.95 | 0.95 | 0.955 | #N/A | #N/A | #N/A |
| 34 | 03.med | conex.full | 03 | med | no | 2 | 1 | 6 | 849 | 15 | 349 | 0.429 | 0.276 | 100 | 0.007 | 0.014 | 0.017 | #N/A | #N/A | #N/A | #N/A | #N/A | 0.144 | 0.918 | 0.909 | 0.923 | 0.96 | 0.954 | 0.963 | #N/A | #N/A | #N/A |
| 35 | 03.sl | conex.full | 03 | sl | no | 1 | 1 | 4 | 694 | 17 | 348 | 0.589 | 0.213 | 100 | 0.006 | 0.015 | 0.023 | #N/A | #N/A | #N/A | #N/A | #N/A | #N/A | 0.944 | 0.942 | 0.948 | 0.969 | 0.966 | 0.973 | #N/A | #N/A | #N/A |
| 36 | 04.ac | conex.full | 04 | ac | yes | 7 | 2 | 17 | 305 | 24 | 49 | 0.576 | 0.102 | 94 | 0.017 | 0.034 | 0.024 | 0.962 | 0.878 | 0.958 | #N/A | 0.931 | 0.317 | 0.948 | 0.943 | 0.953 | 0.901 | 0.894 | 0.908 | 1 | 0.574 | 1 |
| 37 | 04.len | conex.full | 04 | len | no | 2 | 1 | 6 | 213 | 13 | 102 | 0.874 | 0.07 | 100 | 0.01 | 0.027 | 0.04 | #N/A | #N/A | #N/A | #N/A | #N/A | #N/A | 0.98 | 0.973 | 0.984 | 0.985 | 0.978 | 0.986 | #N/A | #N/A | #N/A |
| 38 | 04.lenc | conex.full | 04 | lenc | no | 2 | 1 | 7 | 726 | 27 | 301 | 0.451 | 0.233 | 98 | 0.008 | 0.023 | 0.018 | #N/A | #N/A | #N/A | #N/A | #N/A | #N/A | 0.87 | 0.868 | 0.873 | 0.932 | 0.921 | 0.938 | #N/A | #N/A | #N/A |
| 39 | 04.med | conex.full | 04 | med | no | 2 | 1 | 8.5 | 567 | 18 | 194 | 0.497 | 0.18 | 100 | 0.009 | 0.017 | 0.02 | #N/A | #N/A | #N/A | #N/A | #N/A | 0.335 | 0.957 | 0.952 | 0.959 | 0.942 | 0.938 | 0.943 | #N/A | #N/A | #N/A |
| 40 | 04.sl | conex.full | 04 | sl | no | 2 | 1 | 6 | 459 | 22 | 198 | 0.689 | 0.151 | 100 | 0.008 | 0.017 | 0.029 | #N/A | #N/A | #N/A | #N/A | #N/A | #N/A | 0.961 | 0.956 | 0.962 | 0.97 | 0.967 | 0.973 | #N/A | #N/A | #N/A |
| 41 | 05.ac | conex.full | 05 | ac | yes | 9 | 3 | 22 | 204 | 27 | 26 | 0.711 | 0.064 | 95 | 0.019 | 0.036 | 0.03 | 0.986 | 0.936 | 0.963 | #N/A | 0.864 | 0.492 | 0.969 | 0.96 | 0.97 | 0.902 | 0.894 | 0.913 | 1 | 0.658 | 1 |
| 42 | 05.len | conex.full | 05 | len | no | 2 | 1 | 8 | 147 | 12 | 63 | 0.896 | 0.052 | 100 | 0.012 | 0.036 | 0.047 | #N/A | #N/A | #N/A | #N/A | #N/A | #N/A | 0.984 | 0.979 | 0.986 | 0.965 | 0.956 | 0.967 | #N/A | #N/A | #N/A |
| 43 | 05.lenc | conex.full | 05 | lenc | no | 3 | 1 | 9 | 499 | 27 | 188 | 0.561 | 0.158 | 99 | 0.01 | 0.028 | 0.021 | #N/A | #N/A | #N/A | #N/A | #N/A | #N/A | 0.896 | 0.887 | 0.9 | 0.924 | 0.908 | 0.931 | #N/A | #N/A | #N/A |
| 44 | 05.med | conex.full | 05 | med | no | 3 | 1 | 12 | 372 | 20 | 106 | 0.609 | 0.119 | 100 | 0.012 | 0.021 | 0.026 | #N/A | #N/A | #N/A | #N/A | 0.87 | 0.420 | 0.97 | 0.964 | 0.974 | 0.947 | 0.935 | 0.957 | 1 | 1 | 1 |
| 45 | 05.sl | conex.full | 05 | sl | no | 2 | 1 | 7 | 320 | 20 | 139 | 0.773 | 0.102 | 100 | 0.01 | 0.021 | 0.035 | #N/A | #N/A | #N/A | #N/A | #N/A | #N/A | 0.97 | 0.96 | 0.976 | 0.973 | 0.966 | 0.976 | #N/A | #N/A | #N/A |
| 46 | 06.ac | conex.full | 06 | ac | yes | 13 | 4 | 41 | 149 | 30 | 17 | 0.779 | 0.041 | 91 | 0.022 | 0.043 | 0.034 | 0.988 | 0.937 | 0.974 | #N/A | 0.871 | 0.853 | 0.962 | 0.96 | 0.966 | 0.862 | 0.838 | 0.882 | 1 | 0.638 | 1 |
| 47 | 07.ac | conex.full | 07 | ac | yes | 16 | 5 | 52.5 | 119 | 31 | 12 | 0.816 | 0.029 | 90 | 0.023 | 0.044 | 0.037 | 0.991 | 0.972 | 0.973 | #N/A | 0.861 | 0.945 | 0.96 | 0.958 | 0.968 | 0.847 | 0.815 | 0.859 | 1 | 0.628 | 1 |
| 48 | 08.ac | conex.full | 08 | ac | yes | 20 | 7 | 64.5 | 100 | 29 | 9 | 0.835 | 0.023 | 84 | 0.025 | 0.048 | 0.039 | 0.991 | 0.971 | 0.975 | #N/A | 0.862 | 0.938 | 0.965 | 0.962 | 0.968 | 0.787 | 0.769 | 0.824 | 1 | 0.288 | 1 |
| 49 | 02.ac | conex.dedup | 02 | ac | no | 3 | 1 | 7 | 985 | 10 | 247 | 0.174 | 0.416 | 91 | 0.01 | 0.027 | 0.012 | #N/A | #N/A | #N/A | #N/A | #N/A | #N/A | 0.841 | 0.823 | 0.849 | 0.946 | 0.943 | 0.951 | #N/A | #N/A | #N/A |
| 50 | 02.len | conex.dedup | 02 | len | no | 1 | 1 | 3 | 675 | 14 | 379 | 0.663 | 0.194 | 100 | 0.006 | 0.015 | 0.022 | #N/A | #N/A | #N/A | #N/A | #N/A | #N/A | 0.92 | 0.916 | 0.927 | 0.969 | 0.966 | 0.971 | #N/A | #N/A | #N/A |
| 51 | 02.lenc | conex.dedup | 02 | lenc | no | 2 | 1 | 3 | 1681 | 12 | 826 | 0.172 | 0.536 | 99 | 0.004 | 0.012 | 0.012 | #N/A | #N/A | #N/A | #N/A | #N/A | #N/A | 0.839 | 0.827 | 0.847 | 0.965 | 0.96 | 0.973 | #N/A | #N/A | #N/A |
| 52 | 02.med | conex.dedup | 02 | med | no | 1 | 1 | 3 | 1792 | 7 | 916 | 0.157 | 0.572 | 99 | 0.004 | 0.008 | 0.011 | #N/A | #N/A | #N/A | #N/A | #N/A | #N/A | 0.851 | 0.835 | 0.854 | 0.973 | 0.967 | 0.978 | #N/A | #N/A | #N/A |
| 53 | 02.sl | conex.dedup | 02 | sl | no | 1 | 1 | 3 | 1098 | 14 | 602 | 0.464 | 0.314 | 100 | 0.004 | 0.01 | 0.017 | #N/A | #N/A | #N/A | #N/A | #N/A | #N/A | 0.919 | 0.904 | 0.921 | 0.969 | 0.963 | 0.972 | #N/A | #N/A | #N/A |
| 54 | 03.ac | conex.dedup | 03 | ac | no | 4 | 2 | 11 | 541 | 14 | 94 | 0.383 | 0.223 | 93 | 0.013 | 0.03 | 0.017 | #N/A | #N/A | #N/A | #N/A | #N/A | 0.162 | 0.925 | 0.918 | 0.931 | 0.928 | 0.921 | 0.935 | #N/A | #N/A | #N/A |
| 55 | 03.len | conex.dedup | 03 | len | no | 1 | 1 | 4.25 | 360 | 15 | 184 | 0.803 | 0.111 | 100 | 0.008 | 0.021 | 0.031 | #N/A | #N/A | #N/A | #N/A | #N/A | #N/A | 0.957 | 0.954 | 0.961 | 0.973 | 0.97 | 0.978 | #N/A | #N/A | #N/A |
| 56 | 03.lenc | conex.dedup | 03 | lenc | no | 2 | 1 | 5 | 1038 | 18 | 437 | 0.286 | 0.348 | 98 | 0.006 | 0.017 | 0.015 | #N/A | #N/A | #N/A | #N/A | #N/A | #N/A | 0.863 | 0.853 | 0.867 | 0.948 | 0.945 | 0.956 | #N/A | #N/A | #N/A |
| 57 | 03.med | conex.dedup | 03 | med | no | 2 | 1 | 5 | 914 | 13 | 381 | 0.401 | 0.312 | 100 | 0.006 | 0.013 | 0.016 | #N/A | #N/A | #N/A | #N/A | #N/A | 0.153 | 0.937 | 0.926 | 0.948 | 0.957 | 0.955 | 0.962 | #N/A | #N/A | #N/A |
| 58 | 03.sl | conex.dedup | 03 | sl | no | 2 | 1 | 4 | 660 | 20 | 323 | 0.618 | 0.206 | 100 | 0.007 | 0.015 | 0.023 | #N/A | #N/A | #N/A | #N/A | #N/A | #N/A | 0.952 | 0.95 | 0.954 | 0.968 | 0.967 | 0.97 | #N/A | #N/A | #N/A |
| 59 | 04.ac | conex.dedup | 04 | ac | yes | 6 | 2 | 17 | 324 | 24 | 41 | 0.572 | 0.116 | 96 | 0.016 | 0.036 | 0.023 | 0.95 | 0.872 | 0.953 | #N/A | 0.932 | 0.353 | 0.951 | 0.944 | 0.966 | 0.927 | 0.916 | 0.93 | 1 | 0.713 | 1 |
| 60 | 04.len | conex.dedup | 04 | len | no | 2 | 1 | 6 | 211 | 15 | 98 | 0.873 | 0.069 | 100 | 0.011 | 0.029 | 0.04 | #N/A | #N/A | #N/A | #N/A | #N/A | #N/A | 0.979 | 0.974 | 0.982 | 0.983 | 0.978 | 0.984 | #N/A | #N/A | #N/A |
| 61 | 04.lenc | conex.dedup | 04 | lenc | no | 2 | 1 | 7.75 | 682 | 28 | 247 | 0.45 | 0.231 | 96 | 0.009 | 0.023 | 0.019 | #N/A | #N/A | #N/A | #N/A | #N/A | #N/A | 0.869 | 0.864 | 0.875 | 0.931 | 0.922 | 0.937 | #N/A | #N/A | #N/A |
| 62 | 04.med | conex.dedup | 04 | med | no | 2 | 1 | 7 | 613 | 19 | 215 | 0.505 | 0.218 | 100 | 0.009 | 0.017 | 0.021 | #N/A | #N/A | #N/A | #N/A | #N/A | 0.358 | 0.961 | 0.95 | 0.972 | 0.956 | 0.954 | 0.966 | #N/A | #N/A | #N/A |
| 63 | 04.sl | conex.dedup | 04 | sl | no | 2 | 1 | 6 | 463 | 22 | 198 | 0.682 | 0.153 | 100 | 0.009 | 0.018 | 0.029 | #N/A | #N/A | #N/A | #N/A | #N/A | #N/A | 0.96 | 0.958 | 0.962 | 0.969 | 0.966 | 0.972 | #N/A | #N/A | #N/A |
| 64 | 05.ac | conex.dedup | 05 | ac | yes | 8.5 | 3 | 21.25 | 240 | 26 | 26 | 0.645 | 0.073 | 94 | 0.018 | 0.04 | 0.026 | 0.95 | 0.867 | 0.96 | #N/A | 0.793 | 0.426 | 0.951 | 0.946 | 0.962 | 0.907 | 0.881 | 0.91 | 1 | 0.728 | 1 |
| 65 | 05.len | conex.dedup | 05 | len | no | 2 | 1 | 8 | 141 | 12 | 60 | 0.901 | 0.051 | 100 | 0.013 | 0.04 | 0.048 | #N/A | #N/A | #N/A | #N/A | #N/A | #N/A | 0.983 | 0.98 | 0.986 | 0.959 | 0.956 | 0.967 | #N/A | #N/A | #N/A |
| 66 | 05.lenc | conex.dedup | 05 | lenc | no | 3 | 1 | 10 | 481 | 34 | 148 | 0.562 | 0.161 | 96.5 | 0.01 | 0.028 | 0.021 | #N/A | #N/A | #N/A | #N/A | #N/A | #N/A | 0.877 | 0.871 | 0.879 | 0.898 | 0.888 | 0.91 | #N/A | #N/A | #N/A |
| 67 | 05.med | conex.dedup | 05 | med | no | 3 | 1 | 10 | 427 | 23 | 116 | 0.574 | 0.156 | 100 | 0.011 | 0.02 | 0.024 | 0.947 | 0.859 | #N/A | #N/A | 0.899 | 0.464 | 0.962 | 0.954 | 0.966 | 0.945 | 0.94 | 0.949 | 1 | 1 | 1 |
| 68 | 05.sl | conex.dedup | 05 | sl | no | 2 | 1 | 8 | 321 | 21 | 139 | 0.767 | 0.102 | 100 | 0.011 | 0.021 | 0.035 | #N/A | #N/A | #N/A | #N/A | #N/A | #N/A | 0.974 | 0.969 | 0.976 | 0.972 | 0.97 | 0.976 | #N/A | #N/A | #N/A |
| 69 | 06.ac | conex.dedup | 06 | ac | yes | 10.5 | 3 | 25 | 192 | 29 | 20 | 0.721 | 0.054 | 94 | 0.02 | 0.046 | 0.028 | 0.958 | 0.86 | 0.976 | #N/A | 0.803 | 0.875 | 0.95 | 0.944 | 0.962 | 0.863 | 0.854 | 0.885 | 1 | 0.64 | 1 |
| 70 | 07.ac | conex.dedup | 07 | ac | yes | 12.5 | 3 | 39 | 154 | 30 | 19 | 0.775 | 0.04 | 91 | 0.021 | 0.047 | 0.029 | 0.959 | 0.846 | 0.968 | #N/A | 0.792 | 0.949 | 0.948 | 0.945 | 0.958 | 0.815 | 0.804 | 0.839 | 1 | 0.523 | 1 |
| 71 | 08.ac | conex.dedup | 08 | ac | yes | 15 | 4 | 41 | 133 | 28 | 14 | 0.791 | 0.03 | 91 | 0.023 | 0.053 | 0.032 | 0.971 | 0.905 | 0.971 | #N/A | 0.804 | 0.942 | 0.951 | 0.945 | 0.962 | 0.808 | 0.788 | 0.823 | 1 | 0.469 | 1 |
| 72 | 02.ac | ml.dedup | 02 | ac | no | 3 | 1 | 7 | 988 | 13 | 253 | 0.173 | 0.402 | 83 | 0.01 | 0.027 | 0.012 | #N/A | #N/A | #N/A | #N/A | #N/A | #N/A | 0.809 | 0.798 | 0.821 | 0.935 | 0.932 | 0.939 | #N/A | #N/A | #N/A |
| 73 | 02.len | ml.dedup | 02 | len | no | 1 | 1 | 3 | 675 | 15 | 376 | 0.655 | 0.192 | 100 | 0.006 | 0.015 | 0.022 | #N/A | #N/A | #N/A | #N/A | #N/A | #N/A | 0.924 | 0.918 | 0.929 | 0.967 | 0.965 | 0.97 | #N/A | #N/A | #N/A |
| 74 | 02.lenc | ml.dedup | 02 | lenc | no | 2 | 1 | 4 | 1642 | 10 | 814 | 0.161 | 0.529 | 98 | 0.004 | 0.012 | 0.012 | #N/A | #N/A | #N/A | #N/A | #N/A | #N/A | 0.833 | 0.823 | 0.842 | 0.962 | 0.96 | 0.963 | #N/A | #N/A | #N/A |
| 75 | 02.med | ml.dedup | 02 | med | no | 1 | 1 | 3 | 1798 | 9 | 945 | 0.151 | 0.554 | 99 | 0.004 | 0.009 | 0.011 | #N/A | #N/A | #N/A | #N/A | #N/A | #N/A | 0.806 | 0.8 | 0.821 | 0.966 | 0.964 | 0.969 | #N/A | #N/A | #N/A |
| 76 | 02.sl | ml.dedup | 02 | sl | no | 1 | 1 | 3 | 1102 | 12 | 613 | 0.464 | 0.316 | 100 | 0.004 | 0.01 | 0.017 | #N/A | #N/A | #N/A | #N/A | #N/A | #N/A | 0.915 | 0.899 | 0.917 | 0.969 | 0.966 | 0.972 | #N/A | #N/A | #N/A |
| 77 | 03.ac | ml.dedup | 03 | ac | no | 4 | 2 | 11 | 557 | 20 | 119 | 0.39 | 0.225 | 93 | 0.013 | 0.03 | 0.018 | #N/A | #N/A | #N/A | #N/A | #N/A | 0.163 | 0.869 | 0.858 | 0.872 | 0.939 | 0.925 | 0.941 | #N/A | #N/A | #N/A |
| 78 | 03.len | ml.dedup | 03 | len | no | 1 | 1 | 4.25 | 356 | 14 | 184 | 0.803 | 0.111 | 100 | 0.008 | 0.021 | 0.031 | #N/A | #N/A | #N/A | #N/A | #N/A | #N/A | 0.96 | 0.954 | 0.965 | 0.974 | 0.971 | 0.98 | #N/A | #N/A | #N/A |
| 79 | 03.lenc | ml.dedup | 03 | lenc | no | 2 | 1 | 5 | 974 | 19 | 417 | 0.307 | 0.323 | 98 | 0.007 | 0.018 | 0.016 | #N/A | #N/A | #N/A | #N/A | #N/A | #N/A | 0.837 | 0.833 | 0.859 | 0.937 | 0.937 | 0.939 | #N/A | #N/A | #N/A |
| 80 | 03.med | ml.dedup | 03 | med | no | 2 | 1 | 5 | 895 | 13 | 371 | 0.42 | 0.305 | 100 | 0.006 | 0.013 | 0.017 | #N/A | #N/A | #N/A | #N/A | #N/A | 0.152 | 0.942 | 0.939 | 0.95 | 0.962 | 0.957 | 0.965 | #N/A | #N/A | #N/A |
| 81 | 03.sl | ml.dedup | 03 | sl | no | 2 | 1 | 4 | 654 | 18 | 321 | 0.611 | 0.202 | 100 | 0.007 | 0.015 | 0.024 | #N/A | #N/A | #N/A | #N/A | #N/A | #N/A | 0.952 | 0.947 | 0.957 | 0.972 | 0.967 | 0.975 | #N/A | #N/A | #N/A |
| 82 | 04.ac | ml.dedup | 04 | ac | yes | 7 | 2 | 17 | 323 | 22 | 52 | 0.548 | 0.109 | 96 | 0.017 | 0.038 | 0.024 | 0.974 | 0.91 | 0.934 | #N/A | 0.972 | 0.348 | 0.944 | 0.94 | 0.948 | 0.909 | 0.897 | 0.91 | 1 | 0.586 | 1 |
| 83 | 04.len | ml.dedup | 04 | len | no | 2 | 1 | 6 | 206 | 12 | 97 | 0.88 | 0.069 | 100 | 0.01 | 0.029 | 0.041 | #N/A | #N/A | #N/A | #N/A | #N/A | #N/A | 0.98 | 0.974 | 0.984 | 0.986 | 0.978 | 0.986 | #N/A | #N/A | #N/A |
| 84 | 04.lenc | ml.dedup | 04 | lenc | no | 2 | 1 | 8 | 596 | 30 | 231 | 0.515 | 0.19 | 98 | 0.009 | 0.024 | 0.021 | #N/A | #N/A | #N/A | #N/A | #N/A | #N/A | 0.862 | 0.853 | 0.872 | 0.924 | 0.912 | 0.93 | #N/A | #N/A | #N/A |
| 85 | 04.med | ml.dedup | 04 | med | no | 2 | 1 | 7 | 632 | 18 | 231 | 0.487 | 0.214 | 100 | 0.009 | 0.017 | 0.02 | #N/A | #N/A | #N/A | #N/A | #N/A | 0.344 | 0.952 | 0.95 | 0.954 | 0.945 | 0.939 | 0.95 | #N/A | #N/A | #N/A |
| 86 | 04.sl | ml.dedup | 04 | sl | no | 2 | 1 | 6 | 461 | 21 | 199 | 0.683 | 0.153 | 100 | 0.009 | 0.018 | 0.029 | #N/A | #N/A | #N/A | #N/A | #N/A | #N/A | 0.959 | 0.954 | 0.964 | 0.969 | 0.967 | 0.976 | #N/A | #N/A | #N/A |
| 87 | 05.ac | ml.dedup | 05 | ac | yes | 9 | 3 | 21 | 216 | 26 | 30 | 0.693 | 0.067 | 97 | 0.02 | 0.041 | 0.03 | 0.991 | 0.935 | 0.957 | #N/A | 0.779 | 0.500 | 0.969 | 0.966 | 0.971 | 0.909 | 0.9 | 0.919 | 1 | 0.598 | 1 |
| 88 | 05.len | ml.dedup | 05 | len | no | 2 | 1 | 8 | 138 | 11 | 57 | 0.904 | 0.05 | 100 | 0.014 | 0.041 | 0.049 | #N/A | #N/A | #N/A | #N/A | #N/A | #N/A | 0.982 | 0.981 | 0.988 | 0.973 | 0.965 | 0.976 | #N/A | #N/A | #N/A |
| 89 | 05.lenc | ml.dedup | 05 | lenc | no | 3 | 1 | 10 | 397 | 33 | 136 | 0.674 | 0.128 | 98 | 0.011 | 0.03 | 0.025 | #N/A | #N/A | #N/A | #N/A | #N/A | #N/A | 0.89 | 0.887 | 0.895 | 0.917 | 0.91 | 0.92 | #N/A | #N/A | #N/A |
| 90 | 05.med | ml.dedup | 05 | med | no | 3 | 1 | 11 | 384 | 20 | 119 | 0.606 | 0.132 | 100 | 0.012 | 0.022 | 0.026 | 0.992 | 0.968 | #N/A | #N/A | 0.885 | 0.465 | 0.97 | 0.968 | 0.971 | 0.941 | 0.937 | 0.95 | 1 | 1 | 1 |
| 91 | 05.sl | ml.dedup | 05 | sl | no | 2 | 1 | 8 | 323 | 21 | 136 | 0.756 | 0.104 | 100 | 0.011 | 0.021 | 0.035 | #N/A | #N/A | #N/A | #N/A | #N/A | #N/A | 0.97 | 0.966 | 0.974 | 0.973 | 0.969 | 0.975 | #N/A | #N/A | #N/A |
| 92 | 06.ac | ml.dedup | 06 | ac | yes | 11 | 3 | 29 | 177 | 28 | 28 | 0.737 | 0.047 | 95.5 | 0.021 | 0.043 | 0.032 | 0.991 | 0.944 | 0.954 | #N/A | 0.781 | 0.883 | 0.968 | 0.963 | 0.968 | 0.878 | 0.87 | 0.884 | 1 | 0.6 | 1 |
| 93 | 07.ac | ml.dedup | 07 | ac | yes | 11 | 3.5 | 41 | 139 | 29 | 23 | 0.808 | 0.036 | 90.5 | 0.023 | 0.044 | 0.035 | 0.992 | 0.955 | 0.965 | #N/A | 0.874 | 0.936 | 0.967 | 0.964 | 0.972 | 0.859 | 0.845 | 0.872 | 1 | 0.771 | 1 |
| 94 | 08.ac | ml.dedup | 08 | ac | yes | 13 | 2 | 43 | 114 | 28 | 21 | 0.852 | 0.026 | 91.5 | 0.025 | 0.045 | 0.037 | 0.997 | 0.995 | 0.957 | #N/A | 0.84 | 0.951 | 0.967 | 0.963 | 0.969 | 0.828 | 0.82 | 0.847 | 1 | 0.792 | 1 |
| 95 | 02.ac | con50.dedup | 02 | ac | no | 4 | 2 | 8 | 815 | 17 | 173 | 0.304 | 0.345 | 94 | 0.011 | 0.028 | 0.014 | #N/A | #N/A | #N/A | #N/A | #N/A | #N/A | 0.852 | 0.847 | 0.859 | 0.952 | 0.945 | 0.955 | #N/A | #N/A | #N/A |
| 96 | 02.len | con50.dedup | 02 | len | no | 1 | 1 | 3 | 699 | 17 | 386 | 0.635 | 0.2 | 100 | 0.006 | 0.017 | 0.022 | #N/A | #N/A | #N/A | #N/A | #N/A | #N/A | 0.919 | 0.913 | 0.92 | 0.963 | 0.959 | 0.965 | #N/A | #N/A | #N/A |
| 97 | 02.lenc | con50.dedup | 02 | lenc | no | 2 | 1 | 3 | 1806 | 7 | 887 | 0.105 | 0.588 | 99 | 0.004 | 0.012 | 0.011 | #N/A | #N/A | #N/A | #N/A | #N/A | #N/A | 0.866 | 0.861 | 0.875 | 0.974 | 0.969 | 0.976 | #N/A | #N/A | #N/A |
| 98 | 02.med | con50.dedup | 02 | med | no | 1 | 1 | 3 | 1714 | 12 | 883 | 0.196 | 0.542 | 99.5 | 0.004 | 0.008 | 0.012 | #N/A | #N/A | #N/A | #N/A | #N/A | #N/A | 0.816 | 0.81 | 0.825 | 0.971 | 0.968 | 0.973 | #N/A | #N/A | #N/A |
| 99 | 02.sl | con50.dedup | 02 | sl | no | 1 | 1 | 3 | 1114 | 12 | 615 | 0.46 | 0.321 | 100 | 0.004 | 0.01 | 0.017 | #N/A | #N/A | #N/A | #N/A | #N/A | #N/A | 0.909 | 0.903 | 0.915 | 0.969 | 0.967 | 0.972 | #N/A | #N/A | #N/A |
| 100 | 03.ac | con50.dedup | 03 | ac | no | 5 | 2 | 12 | 428 | 16 | 72 | 0.487 | 0.171 | 98 | 0.015 | 0.034 | 0.022 | #N/A | #N/A | #N/A | #N/A | #N/A | 0.163 | 0.963 | 0.954 | 0.97 | 0.95 | 0.942 | 0.953 | #N/A | #N/A | #N/A |
| 101 | 03.len | con50.dedup | 03 | len | no | 1 | 1 | 4 | 382 | 14 | 195 | 0.779 | 0.117 | 100 | 0.008 | 0.021 | 0.03 | #N/A | #N/A | #N/A | #N/A | #N/A | #N/A | 0.956 | 0.954 | 0.959 | 0.972 | 0.971 | 0.979 | #N/A | #N/A | #N/A |
| 102 | 03.lenc | con50.dedup | 03 | lenc | no | 2 | 1 | 5 | 1117 | 17 | 470 | 0.241 | 0.381 | 98 | 0.006 | 0.017 | 0.015 | #N/A | #N/A | #N/A | #N/A | #N/A | #N/A | 0.883 | 0.866 | 0.899 | 0.957 | 0.949 | 0.964 | #N/A | #N/A | #N/A |
| 103 | 03.med | con50.dedup | 03 | med | no | 2 | 1 | 5 | 846 | 12 | 340 | 0.428 | 0.287 | 100 | 0.006 | 0.013 | 0.017 | #N/A | #N/A | #N/A | #N/A | #N/A | 0.154 | 0.948 | 0.943 | 0.958 | 0.963 | 0.956 | 0.966 | #N/A | #N/A | #N/A |
| 104 | 03.sl | con50.dedup | 03 | sl | no | 2 | 1 | 4 | 671 | 19 | 331 | 0.608 | 0.208 | 100 | 0.007 | 0.015 | 0.023 | #N/A | #N/A | #N/A | #N/A | #N/A | #N/A | 0.949 | 0.946 | 0.952 | 0.971 | 0.967 | 0.973 | #N/A | #N/A | #N/A |
| 105 | 04.ac | con50.dedup | 04 | ac | yes | 8 | 3 | 17 | 284 | 23 | 33 | 0.599 | 0.104 | 98 | 0.018 | 0.039 | 0.026 | 0.998 | 0.985 | 0.959 | #N/A | 0.909 | 0.330 | 0.968 | 0.965 | 0.971 | 0.943 | 0.931 | 0.949 | 1 | 0.731 | 1 |
| 106 | 04.len | con50.dedup | 04 | len | no | 2 | 1 | 8 | 225 | 15 | 100 | 0.851 | 0.077 | 100 | 0.011 | 0.029 | 0.04 | #N/A | #N/A | #N/A | #N/A | #N/A | #N/A | 0.977 | 0.972 | 0.98 | 0.983 | 0.975 | 0.985 | #N/A | #N/A | #N/A |
| 107 | 04.lenc | con50.dedup | 04 | lenc | no | 2 | 1 | 7 | 769 | 28 | 287 | 0.396 | 0.262 | 97 | 0.008 | 0.022 | 0.017 | #N/A | #N/A | #N/A | #N/A | #N/A | #N/A | 0.884 | 0.873 | 0.899 | 0.939 | 0.93 | 0.942 | #N/A | #N/A | #N/A |
| 108 | 04.med | con50.dedup | 04 | med | no | 2 | 1 | 7.75 | 586 | 18 | 213 | 0.532 | 0.201 | 100 | 0.009 | 0.017 | 0.022 | #N/A | #N/A | #N/A | #N/A | #N/A | 0.364 | 0.974 | 0.97 | 0.98 | 0.971 | 0.964 | 0.972 | #N/A | #N/A | #N/A |
| 109 | 04.sl | con50.dedup | 04 | sl | no | 2 | 1 | 6 | 471 | 20 | 200 | 0.668 | 0.156 | 100 | 0.009 | 0.018 | 0.029 | #N/A | #N/A | #N/A | #N/A | #N/A | #N/A | 0.962 | 0.959 | 0.967 | 0.97 | 0.968 | 0.978 | #N/A | #N/A | #N/A |
| 110 | 05.ac | con50.dedup | 05 | ac | yes | 9 | 3 | 20 | 219 | 25 | 23 | 0.689 | 0.074 | 98 | 0.02 | 0.041 | 0.03 | 0.994 | 0.951 | 0.966 | #N/A | 0.842 | 0.499 | 0.974 | 0.974 | 0.975 | 0.941 | 0.933 | 0.946 | 1 | 0.783 | 1 |
| 111 | 05.len | con50.dedup | 05 | len | no | 2 | 1 | 8 | 149 | 13 | 62 | 0.893 | 0.051 | 100 | 0.013 | 0.037 | 0.047 | #N/A | #N/A | #N/A | #N/A | #N/A | #N/A | 0.984 | 0.981 | 0.985 | 0.971 | 0.956 | 0.975 | #N/A | #N/A | #N/A |
| 112 | 05.lenc | con50.dedup | 05 | lenc | no | 2 | 1 | 8 | 579 | 32 | 198 | 0.528 | 0.194 | 97 | 0.009 | 0.025 | 0.019 | #N/A | #N/A | #N/A | #N/A | #N/A | #N/A | 0.9 | 0.882 | 0.911 | 0.917 | 0.913 | 0.924 | #N/A | #N/A | #N/A |
| 113 | 05.med | con50.dedup | 05 | med | no | 3 | 1 | 10 | 398 | 21 | 117 | 0.598 | 0.141 | 100 | 0.012 | 0.021 | 0.026 | 0.991 | 0.944 | #N/A | #N/A | 0.941 | 0.462 | 0.979 | 0.972 | 0.98 | 0.965 | 0.952 | 0.967 | 1 | 1 | 1 |
| 114 | 05.sl | con50.dedup | 05 | sl | no | 2 | 1 | 8 | 327 | 22 | 138 | 0.761 | 0.105 | 100 | 0.011 | 0.021 | 0.034 | #N/A | #N/A | #N/A | #N/A | #N/A | #N/A | 0.97 | 0.964 | 0.973 | 0.972 | 0.968 | 0.974 | #N/A | #N/A | #N/A |
| **115** | **06.ac** | **con50.dedup** | **06** | **ac** | **yes** | **11** | **4** | **25** | **181** | **27** | **19** | **0.729** | **0.049** | **98** | **0.021** | **0.043** | **0.032** | **0.994** | **0.961** | **0.965** | **8.7** | **0.841** | **0.876** | **0.977** | **0.974** | **0.978** | **0.925** | **0.921** | **0.935** | **1** | **0.8** | **1** |
| **116** | **07.ac** | **con50.dedup** | **07** | **ac** | **yes** | **11** | **4** | **34** | **151** | **29** | **17** | **0.792** | **0.04** | **96** | **0.023** | **0.044** | **0.033** | **0.992** | **0.938** | **0.976** | **8.7** | **0.831** | **0.947** | **0.973** | **0.971** | **0.979** | **0.897** | **0.89** | **0.913** | **1** | **0.784** | **1** |
| **117** | **08.ac** | **con50.dedup** | **08** | **ac** | **yes** | **14** | **4.5** | **52** | **115** | **30** | **11** | **0.836** | **0.026** | **90.5** | **0.025** | **0.053** | **0.036** | **0.997** | **0.988** | **0.965** | **6.5** | **0.802** | **0.940** | **0.967** | **0.964** | **0.972** | **0.878** | **0.867** | **0.895** | **1** | **0.801** | **1** |
| 118 | 02.ac | con50.full | 02 | ac | no | 3.5 | 2 | 8 | 768 | 10 | 182 | 0.327 | 0.322 | 98 | 0.011 | 0.028 | 0.015 | #N/A | #N/A | #N/A | #N/A | #N/A | #N/A | 0.915 | 0.9 | 0.933 | 0.951 | 0.94 | 0.964 | #N/A | #N/A | #N/A |
| 119 | 02.len | con50.full | 02 | len | no | 1 | 1 | 3 | 690 | 16 | 383 | 0.636 | 0.196 | 100 | 0.006 | 0.017 | 0.022 | #N/A | #N/A | #N/A | #N/A | #N/A | #N/A | 0.927 | 0.925 | 0.931 | 0.966 | 0.962 | 0.967 | #N/A | #N/A | #N/A |
| 120 | 02.lenc | con50.full | 02 | lenc | no | 2 | 1 | 4 | 1645 | 12 | 806 | 0.15 | 0.53 | 100 | 0.004 | 0.012 | 0.012 | #N/A | #N/A | #N/A | #N/A | #N/A | #N/A | 0.829 | 0.827 | 0.845 | 0.97 | 0.962 | 0.971 | #N/A | #N/A | #N/A |
| 121 | 02.med | con50.full | 02 | med | no | 1 | 1 | 3 | 1380 | 6 | 716 | 0.325 | 0.446 | 100 | 0.004 | 0.009 | 0.013 | #N/A | #N/A | #N/A | #N/A | #N/A | #N/A | 0.955 | 0.939 | 0.96 | 0.974 | 0.971 | 0.978 | #N/A | #N/A | #N/A |
| 122 | 02.sl | con50.full | 02 | sl | no | 1 | 1 | 3 | 1095 | 13 | 604 | 0.473 | 0.315 | 100 | 0.004 | 0.01 | 0.017 | #N/A | #N/A | #N/A | #N/A | #N/A | #N/A | 0.915 | 0.906 | 0.921 | 0.969 | 0.966 | 0.973 | #N/A | #N/A | #N/A |
| 123 | 03.ac | con50.full | 03 | ac | no | 6 | 2 | 14 | 389 | 17 | 61 | 0.506 | 0.153 | 97 | 0.015 | 0.034 | 0.021 | #N/A | #N/A | #N/A | #N/A | #N/A | 0.161 | 0.954 | 0.948 | 0.963 | 0.936 | 0.932 | 0.953 | #N/A | #N/A | #N/A |
| 124 | 03.len | con50.full | 03 | len | no | 1 | 1 | 5 | 374 | 14 | 188 | 0.784 | 0.117 | 100 | 0.008 | 0.021 | 0.03 | #N/A | #N/A | #N/A | #N/A | #N/A | #N/A | 0.955 | 0.949 | 0.958 | 0.976 | 0.97 | 0.98 | #N/A | #N/A | #N/A |
| 125 | 03.lenc | con50.full | 03 | lenc | no | 2 | 1 | 6 | 1089 | 18 | 460 | 0.221 | 0.361 | 99 | 0.006 | 0.017 | 0.015 | #N/A | #N/A | #N/A | #N/A | #N/A | #N/A | 0.847 | 0.835 | 0.855 | 0.954 | 0.947 | 0.957 | #N/A | #N/A | #N/A |
| 126 | 03.med | con50.full | 03 | med | no | 2 | 1 | 5.75 | 798 | 13 | 334 | 0.466 | 0.269 | 100 | 0.007 | 0.015 | 0.017 | #N/A | #N/A | #N/A | #N/A | #N/A | 0.154 | 0.962 | 0.96 | 0.966 | 0.969 | 0.967 | 0.977 | #N/A | #N/A | #N/A |
| 127 | 03.sl | con50.full | 03 | sl | no | 2 | 1 | 5 | 666 | 20 | 324 | 0.6 | 0.208 | 100 | 0.006 | 0.015 | 0.023 | #N/A | #N/A | #N/A | #N/A | #N/A | #N/A | 0.948 | 0.947 | 0.953 | 0.969 | 0.967 | 0.972 | #N/A | #N/A | #N/A |
| 128 | 04.ac | con50.full | 04 | ac | yes | 8 | 3 | 19 | 265 | 24 | 25 | 0.612 | 0.094 | 97 | 0.018 | 0.037 | 0.026 | 0.973 | 0.895 | 0.959 | #N/A | 0.924 | 0.339 | 0.966 | 0.961 | 0.969 | 0.928 | 0.923 | 0.933 | 1 | 0.667 | 1 |
| 129 | 04.len | con50.full | 04 | len | no | 2 | 1 | 6 | 217 | 14 | 105 | 0.868 | 0.072 | 100 | 0.01 | 0.027 | 0.039 | #N/A | #N/A | #N/A | #N/A | #N/A | #N/A | 0.98 | 0.974 | 0.982 | 0.982 | 0.973 | 0.984 | #N/A | #N/A | #N/A |
| 130 | 04.lenc | con50.full | 04 | lenc | no | 2 | 1 | 8 | 722 | 30 | 282 | 0.396 | 0.232 | 99 | 0.008 | 0.022 | 0.018 | #N/A | #N/A | #N/A | #N/A | #N/A | #N/A | 0.857 | 0.851 | 0.875 | 0.927 | 0.919 | 0.94 | #N/A | #N/A | #N/A |
| 131 | 04.med | con50.full | 04 | med | no | 2 | 1 | 8 | 535 | 19 | 180 | 0.54 | 0.19 | 100 | 0.009 | 0.017 | 0.022 | #N/A | #N/A | #N/A | #N/A | #N/A | 0.342 | 0.972 | 0.969 | 0.977 | 0.963 | 0.957 | 0.972 | #N/A | #N/A | #N/A |
| 132 | 04.sl | con50.full | 04 | sl | no | 2 | 1 | 6 | 467 | 23 | 196 | 0.672 | 0.153 | 100 | 0.008 | 0.018 | 0.029 | #N/A | #N/A | #N/A | #N/A | #N/A | #N/A | 0.96 | 0.956 | 0.964 | 0.97 | 0.966 | 0.974 | #N/A | #N/A | #N/A |
| 133 | 05.ac | con50.full | 05 | ac | yes | 11 | 4 | 30.75 | 186 | 29 | 16 | 0.708 | 0.055 | 96 | 0.02 | 0.04 | 0.03 | 0.974 | 0.923 | 0.961 | #N/A | 0.915 | 0.504 | 0.963 | 0.956 | 0.964 | 0.886 | 0.871 | 0.892 | 1 | 0.667 | 1 |
| 134 | 05.len | con50.full | 05 | len | no | 2.5 | 1 | 8.25 | 140 | 13 | 60 | 0.9 | 0.048 | 100 | 0.013 | 0.033 | 0.048 | #N/A | #N/A | #N/A | #N/A | #N/A | #N/A | 0.985 | 0.981 | 0.986 | 0.969 | 0.964 | 0.974 | #N/A | #N/A | #N/A |
| 135 | 05.lenc | con50.full | 05 | lenc | no | 3 | 1 | 11 | 441 | 34 | 158 | 0.596 | 0.134 | 99 | 0.011 | 0.029 | 0.022 | #N/A | #N/A | #N/A | #N/A | #N/A | #N/A | 0.887 | 0.869 | 0.891 | 0.892 | 0.882 | 0.904 | #N/A | #N/A | #N/A |
| 136 | 05.med | con50.full | 05 | med | no | 4 | 1 | 13 | 351 | 23 | 90 | 0.621 | 0.115 | 100 | 0.012 | 0.022 | 0.026 | #N/A | #N/A | #N/A | #N/A | 0.907 | 0.428 | 0.969 | 0.964 | 0.972 | 0.945 | 0.937 | 0.946 | 1 | 1 | 1 |
| 137 | 05.sl | con50.full | 05 | sl | no | 2 | 1 | 7 | 319 | 22 | 137 | 0.776 | 0.102 | 100 | 0.01 | 0.021 | 0.035 | #N/A | #N/A | #N/A | #N/A | #N/A | #N/A | 0.972 | 0.968 | 0.976 | 0.968 | 0.963 | 0.97 | #N/A | #N/A | #N/A |
| 138 | 06.ac | con50.full | 06 | ac | yes | 16 | 5 | 46 | 136 | 32 | 10 | 0.781 | 0.032 | 94 | 0.022 | 0.044 | 0.034 | 0.985 | 0.953 | 0.971 | #N/A | 0.906 | 0.861 | 0.956 | 0.948 | 0.959 | 0.817 | 0.806 | 0.836 | 1 | 0.396 | 1 |
| 139 | 07.ac | con50.full | 07 | ac | yes | 17 | 6.5 | 58 | 115 | 32 | 9 | 0.812 | 0.026 | 91.5 | 0.024 | 0.046 | 0.036 | 0.986 | 0.988 | 0.97 | #N/A | 0.885 | 0.938 | 0.957 | 0.95 | 0.958 | 0.796 | 0.786 | 0.798 | 1 | 0.334 | 1 |
| 140 | 08.ac | con50.full | 08 | ac | yes | 16 | 6.5 | 66 | 95 | 32 | 9 | 0.876 | 0.02 | 90.5 | 0.025 | 0.05 | 0.039 | 0.992 | 0.968 | 0.971 | #N/A | 0.814 | 0.936 | 0.964 | 0.957 | 0.965 | 0.788 | 0.769 | 0.789 | 1 | 0.351 | 1 |
