## Supplementary figures and images for "Phylogenetic-based methods for fine-scale classification of PRRSV-2 ORF5 sequences: a comparison of their robustness and reproducibility"

### Supplementary Figure S1

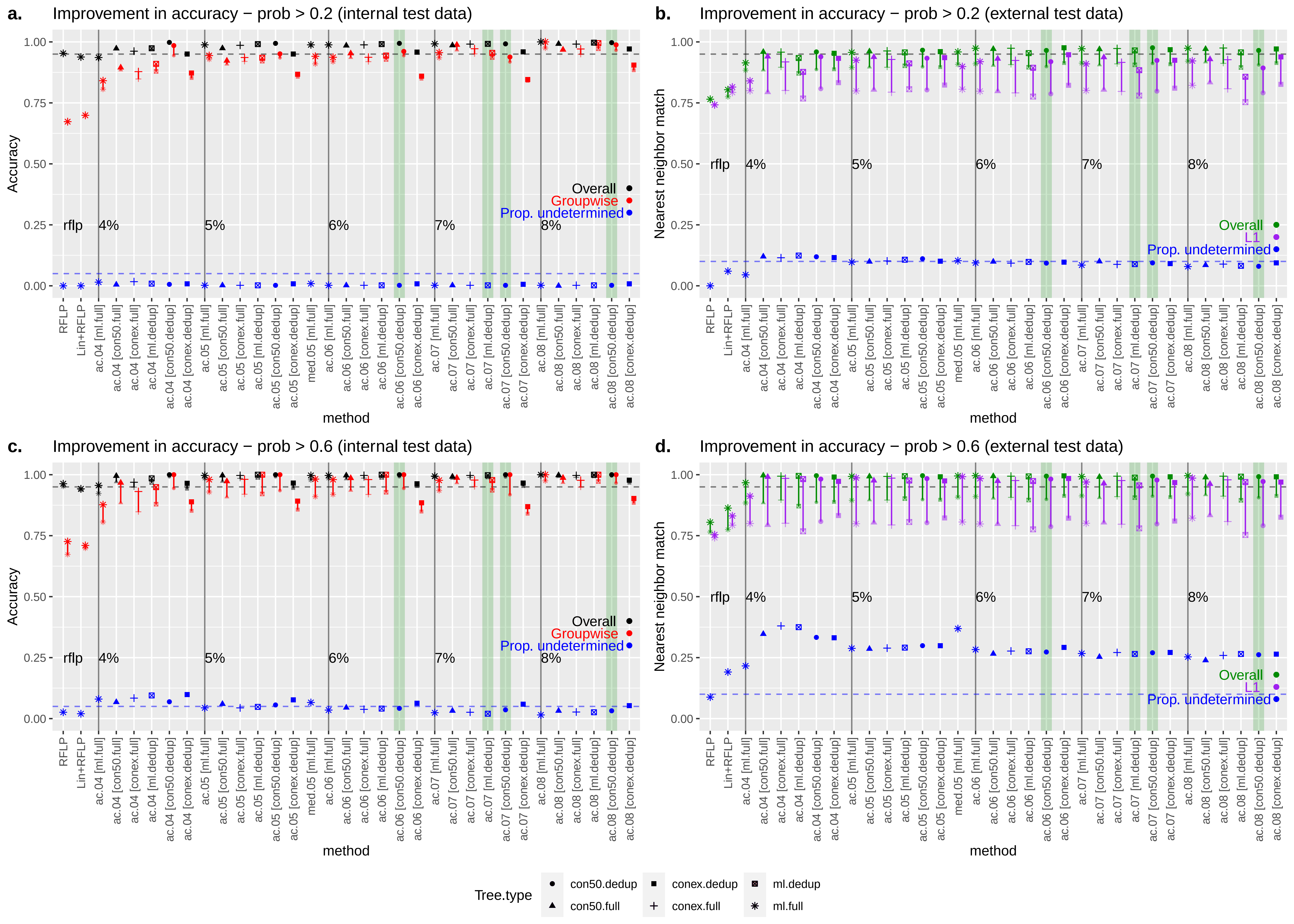

### Supplementary Figure S2

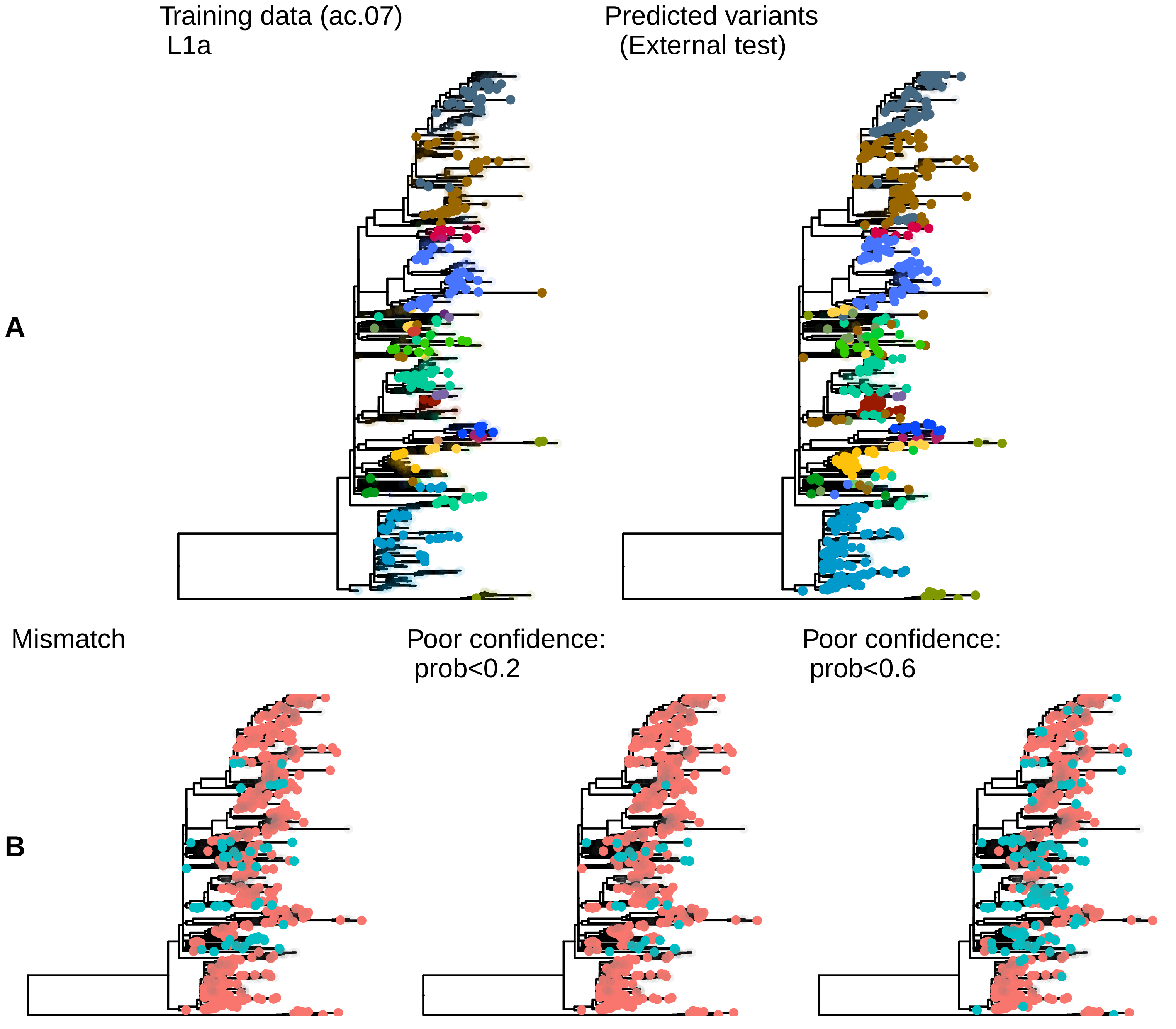

### Supplementary Figure S3

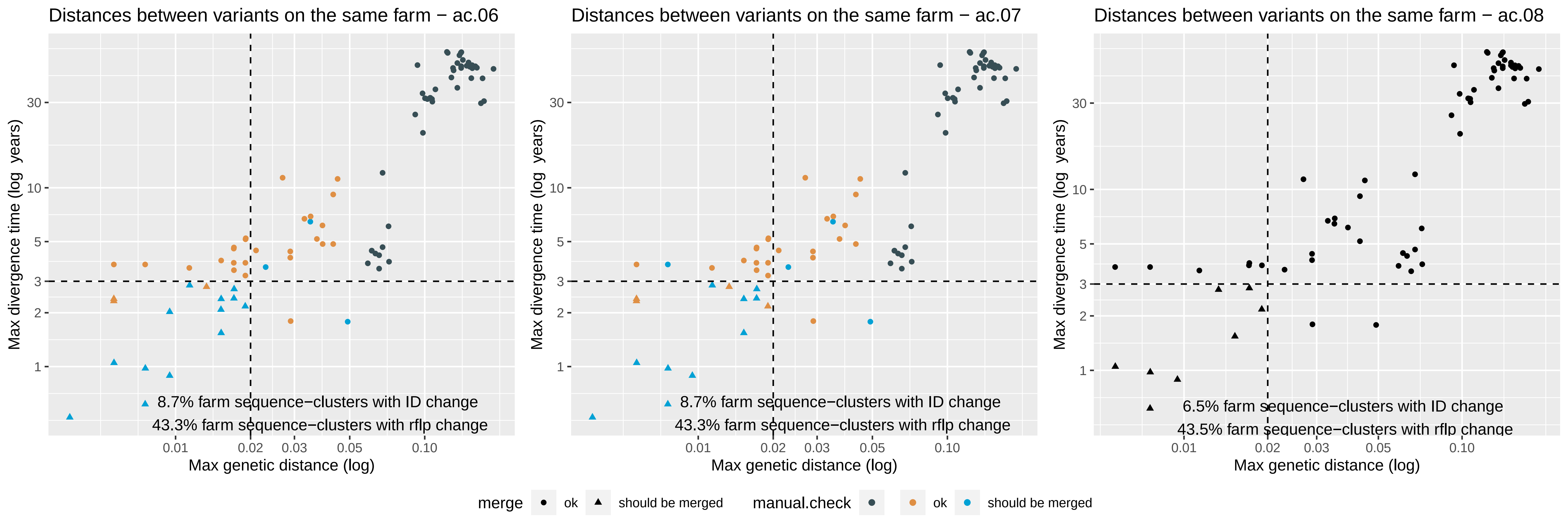
